# A Neural Population Code for Value in Human Orbitofrontal Cortex

**DOI:** 10.64898/2026.03.28.715037

**Authors:** Raphaël Le Bouc, Gilles de Hollander, Marcus Grueschow, Shira M. Lupkin, Vincent B. McGinty, Rafael Polania, Christian C. Ruff

## Abstract

Adaptive behavior depends on the ability to rapidly evaluate options and select those that promise the greatest benefit. Such decisions rely on neural representations of value distributed across multiple brain regions, including the orbitofrontal (OFC) and ventromedial prefrontal cortex (vmPFC), yet the neural code underlying these value representations remains unresolved. The dominant account proposes that OFC/vmPFC neurons encode value through a linear rate code, resulting in a single point estimate at the population level. However, this framework is difficult to reconcile with the heterogeneous tuning observed in individual OFC/vmPFC neurons, which can exhibit both positive and negative correlations with subjective value.

Here, we test the alternative hypothesis—derived from theories of neural coding in perceptual systems— that the OFC/vmPFC implements a probabilistic population code based on non-linear tuning functions. Such tuning allows population activity to represent not only subjective value but also the uncertainty surrounding it, in the form of a flexible posterior probability distribution. Using a population receptive field framework, we fitted non-linear value-tuning functions to functional magnetic resonance imaging data acquired during a value judgment task. Bayesian inversion of this encoding model enabled robust out-of-sample decoding of subjective value across several brain regions, including the OFC/vmPFC. Importantly, value uncertainty estimated from decoded medial OFC/vmPFC posteriors predicted within-subject preference instability, choice stochasticity, and confidence in option values, demonstrating its behavioral relevance and suggesting that participants had conscious access to this information. Complementary single-unit recordings from a subset of monkey OFC neurons similarly revealed nonlinear value-selective tuning.

Together, these findings establish a probabilistic, non-linear population code for value in the OFC/vmPFC. This provides a neural foundation for the probabilistic code through which value, and uncertainty about value, can guide choice.

## INTRODUCTION

Making good decisions —such as selecting the best food source, mate, or habitat— is essential for survival and adaptive fitness. This capacity is thought to depend on brain signals that represent the subjective value of available options and thus guide behaviour in line with the organism’s goals. Such value-related signals have been identified in regions like the orbitofrontal cortex (OFC), ventromedial prefrontal cortex (vmPFC), and the ventral striatum (VS), which together are considered to form a core brain valuation system^1–8^. Understanding the origin and properties of these value signals is critical to elucidating both healthy and pathological goal-directed behaviour.

Bridging the gap between theoretical models of value computation and their underlying neural implementation remains a key challenge. At the computational level, accumulating evidence suggests that value is not passively retrieved from memory but rather actively constructed from different attributes of an option^9–12^. A prominent framework proposes that value construction relies on Bayesian inference, whereby the brain integrates prior beliefs with incoming sensory and contextual evidence to compute a posterior probability distribution over an option’s expected value^13–16^. This probabilistic inference framework offers a parsimonious account of various biases observed in value-based decision-making, such as those induced by choice context^14,15^, the prior distribution of available options^13^, and noise in the internal representations used during inference^13,17^.

At the neural level, however, it remains unclear how neural populations may support such a probabilistic inference of value. A prevailing view suggests that value-related neurons in the OFC/vmPFC employ a linear rate code, where average firing rate correlates with subjective value^18–23^. This idea has shaped dominant population-level analysis approaches, in which reward value is reflected in the mean neuronal activity in the OFC/vmPFC, as measured by Blood-Oxygen-Level-Dependent (BOLD) signal in functional Magnetic Resonance Imaging (fMRI)^1–8^ or by local field potentials (LFP) recorded using intracranial electrodes^20,24,25^. However, the relationship between value and BOLD activity sometimes appears to be reversed in macaques compared to humans^26^, questioning the direct link with a rate code. Moreover, a value coding scheme in which higher firing rate corresponds to higher value is inconsistent with single-unit recordings in the OFC/vmPFC, which reveal highly heterogeneous response profiles. Specifically, only a subset of OFC/vmPFC neurons responds linearly to value^19,20,27–29^, with approximately half of these value-encoding neurons increasing their firing rate with increasing value, while the other half show the opposite pattern^19,20,27–29^, thus predicting no net effect on average population activity. This raises the question how neurons with very different response profiles can collectively encode value signals.

To address this question, we propose a neural coding scheme for subjective value that addresses these shortcomings within a unified framework, and back up this proposal with empirical evidence. Specifically, we build upon theories of probabilistic population coding^30–33^ from perception research. This framework proposes that individual neurons exhibit a non-linear (often bell-shaped) tuning function over stimulus values. As a result, the population activity can collectively encode a flexible posterior probability distribution over the stimulus space. Downstream regions can decode this population response to extract both the most likely stimulus magnitude (mean) and its associated uncertainty (width/precision of the distribution). In perception research, previous studies have provided support for this hypothesis and have linked probabilistic sensory uncertainty, decoded in this way from fMRI brain activity, with perceptual behaviour and choices under uncertainty^34–39^. For example, sensory uncertainty decoded from the visual cortex correlates with errors in the perception of visual orientation^37–39^, motion direction^35^, and visuospatial location^36^. We now propose that the OFC/vmPFC may employ a similar probabilistic population code in which value is represented collectively across neurons that are each tuned to different magnitudes. This coding scheme has several advantages.

First, at the neural level, this framework may explain the variety of observed neuronal response profiles^19,20,27–29^, depending on how offered rewards span different parts of each neuron’s tuning curve. Second, from a theoretical perspective, the coding scheme implies that the activity of these neuronal populations naturally contains information not only about value but also about the uncertainty associated with it. Bayesian decoding approaches formalize how a Bayesian decoder —a downstream brain region or an ideal observer— informed about OFC/vmPFC’s encoding processes could read out from the population activity a probability distribution over value^31,40,41^. A key, testable prediction of this framework is that the uncertainty encoded in the neural population activity should predict preference variability and choice inconsistencies for this probabilistic population code to be functionally meaningful^13^. Furthermore, if individuals have some conscious access to (summary statistics of) the posterior distribution over value, the width of this distribution could serve as a computational substrate for confidence judgments. This aligns with recent theoretical proposals suggesting that confidence in decision-making arises from Bayesian inference—specifically, that reported confidence reflects a summary statistic (e.g., precision or variance) derived from the posterior probability of the decision variable.^42–45^.

Third, such a probabilistic population value code may provide a plausible neural implementation for the Bayesian inference framework for value^13–16^, enabling value computations based on information that is distributed systematically across all neurons in a population. Crucially, if the same principles were at work in multiple neural systems (related to e.g., perception^30–33^, numerical cognition^34,46^, and subjective value), then such a probabilistic coding scheme would go beyond informing theories of OFC/vmPFC function and would suggest how information in sensory and association cortices could be aligned with that in value-related areas like the OFC/vmPFC - regions that share a broadly similar laminar architecture^47^. Thus, the probabilistic population coding scheme would provide a straightforward principle for uncertainty-weighted integration of information across diverse brain systems.

To test whether the human brain represents value using such a probabilistic population code, we analyzed fMRI activity recorded during a preference judgment task. We fitted a population receptive field (pRF) encoding model to the neural data, embedding our assumptions about probabilistic population coding - specifically, non-linear, bell-shaped tuning functions. Next, we inverted the fitted pRF model using Bayesian decoding. This approach allowed us to reliably recover subjective value representations, including both the mean and uncertainty of the value distribution, from value-related brain regions such as the OFC/vmPFC. This approach outperformed decoding based on traditional linear rate-coding models. Beyond value itself, we decoded the precision of neural value representations and found that trial-by-trial estimates of value uncertainty in medial OFC/vmPFC were indeed associated with increased preference variability, greater choice stochasticity, and reduced confidence in option values. To assess the generality of these findings across species and experimental paradigms, we also reanalyzed previously published single-neuron recordings from monkey OFC during an economic decision-making task. This analysis provided convergent evidence for population coding of value, revealing that a subset of OFC neurons exhibits receptive field–like tuning to value. Together, these results establish a unified framework in which the OFC/vmPFC encodes value via a probabilistic population code. This framework accounts for the heterogeneity of OFC/vmPFC neuronal responses, provides a biologically plausible substrate for Bayesian inference over subjective value, explains variability and confidence in decision making through explicit representations of value uncertainty, and suggests a common mechanism by which uncertainty-weighted integration can be shared across perceptual and motivational brain systems.

## RESULTS

### Behavioural signatures of value inference

We aimed to determine whether the human brain uses a non-linear, probabilistic population code to represent value, and whether this scheme enables the readout of both value and its associated uncertainty. If true, this framework would provide a parsimonious, mechanistic explanation for key behavioural signatures of value-based choice. Specifically, imprecision in the probabilistic population code— reflected in greater uncertainty in the decoded posterior distribution—should manifest as increased variability in repeated value judgments, lower reported confidence, and reduced choice accuracy. Because these effects arise from a common source of neural imprecision, they should also correlate across items and individuals. From a purely behavioural perspective, these predictions have already been supported by data demonstrating that a value inference process relying on efficient coding and Bayesian decoding can account for preference variability, confidence, and choice stochasticity^13^. Here, we first replicate these behavioural signatures in an independent dataset, using two complementary behavioural tasks based on a paradigm established in previous work^13^ (Fig. 1a and 1b) for which we examined whether they are consistent with a neural population coding scheme.

**Fig. 1.**
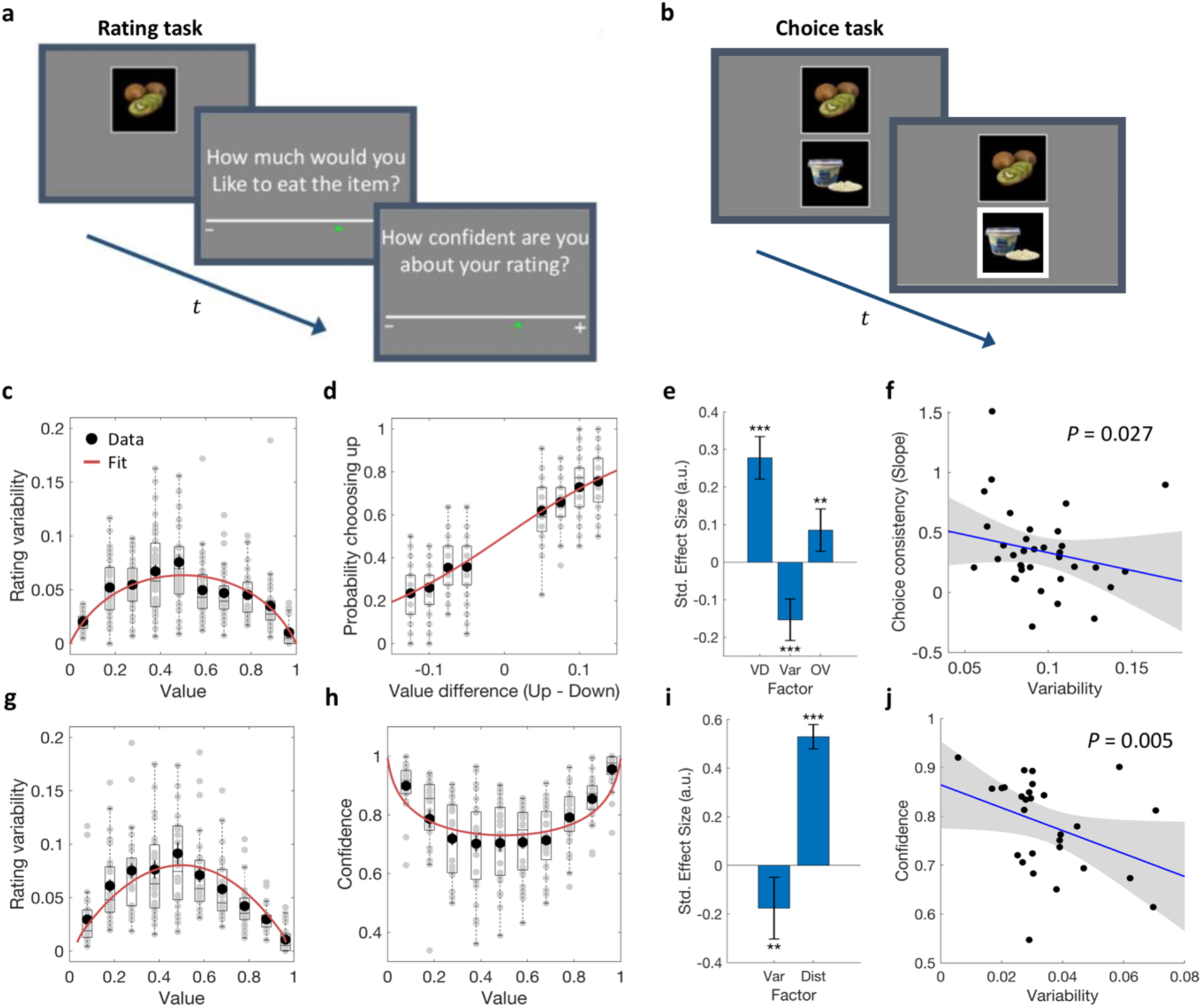
Experimental tasks and behavioural results. **a**, Example display from the rating task (two rounds) during which the participants rated their preference to eat the displayed food item (experiment 1, *n* = 36; experiment 2, *n* = 28) and their confidence in the value ratings (in experiment 2, only) using a continuous rating scale. **b**, Example display from the choice task requiring participants to choose which of the two items they preferred to eat after the experiment. **c**, value rating variability plotted as a function of each item’s mean rating (in 10 value bins) across both rounds for experiment 1. Black dots show the mean across participants; dot error bars represent the s.e.m. across participants; the red curve represents the best fit for a sigmoidal projection mapping of the dispersion of probabilistic representations of value on the rating scale. **d**, Observed choice probability plotted as a function of the two item’s value difference (𝑉_1_ − 𝑉_2_, in 8 bins) for experiment 1. Black dots show the mean across participants; dot error bars represent the s.e.m. across participants; the red curve represents the best sigmoidal fit. **e**, Standardized estimates from multiple logistic regression show that the higher the value difference (VD) between the mean ratings of the choice options, the more consistent the choice. Crucially, the higher the variability (Var) in the rating of the choice options, the less consistent the choice. The overall value of the two options (OV; 𝑉_1_ + 𝑉_2_) was also associated with a higher choice consistency. Error bars represent 95% confidence interval of the posterior estimates. * 𝑃 < 0.05, ** 𝑃 < 0.01, *** 𝑃 < 0.001. **f**, The trial-by-trial effect shown in **e** was also consistent with the negative correlation observed across participants between the global level of variability in the value rating task and the slope of a logistic regression of choice consistency on value difference between options (experiment 1). **g**, Same as **c** but for experiment 2. **h**, confidence ratings plotted as a function of each item’s mean rating (in 10 value bins) averaged across both rounds for experiment 2. Black dots show the mean across participants; dot error bars represent the s.e.m. across participants; the red curve represents the best fit for a sigmoidal projection mapping of the dispersion of probabilistic representations of value on the rating scale. **i**, Standardized estimates from multiple logistic regression show that the higher the rating variability (Var) for a food item, the lower the confidence in that item’s value (experiment 2). Importantly, this controls for the distortion induced by the bounded rating scale, where confidence is higher for items with extreme value ratings (𝐷𝑖𝑠𝑡 = |𝑉 − 0.5|). **j**, This relationship also extends across participants, as reflected by the negative correlation in the rating task in experiment 2, between the global level of variability in value ratings and the mean confidence rating.

In a first behavioural experiment (Experiment 1), 36 participants were asked to provide preference judgments for a set of 64 food items using a continuous value rating scale (Fig. 1a). The task was administered twice consecutively, in order to measure variability in value ratings of the same set of items. Importantly, participants were not told beforehand that a second rating phase would take place. This was done to prevent participants from intentionally memorising the item ratings, thus ensuring a reliable measure of variability in value estimates (Methods, and Fig. 1a). Participants’ value ratings were indeed variable, following an inverted U-shape with a lower variability for extreme values (multiple logistic regression, 𝛽_MixedEffects_ = -0.34 ± 0.03, p < 0.001; Fig. 1c), which may reflect the compression of the value space onto the bounded rating scale^13,48,49^. We then asked the same participants to perform a choice task that consisted in selecting the item they would prefer to eat from pairs of food items they had previously rated (Methods, and Fig. 1b). A choice was considered consistent when participants chose the item they had given the higher average rating across the two rating phases. We first tested whether the variability observed in value ratings (Fig. 1c) would correlate with the variability observed in choices for the same items (Fig. 1d). Replicating previous work^13^, we found that choice consistency was influenced by the value difference between the choice options, with larger differences being associated with higher choice consistency (mixed-effects multiple logistic regression, 𝛽_MixedEffects_ = 0.28 ± 0.03, p < 0.001; Fig. 1e). More importantly, choice consistency was also affected by the variability in the preceding value ratings of the choice items: Choice options that were given more variable ratings initially led to less consistent choices (𝛽_MixedEffects_ = -0.15 ± 0.03, p < 0.001; Fig.1e). Furthermore, this relationship also held across individuals, as participants’ mean level of variability in the value rating task correlated negatively with the slope of the logistic regression of their choices on the mean value difference between choice options (𝛽_Robust_ = –4.33 ± 1.9, p = 0.027; Fig. 1f). Thus, higher variability when estimating the value of options was associated with lower consistency in subsequent choices, both across food items and across individuals. This suggests that both these effects have a common origin in the way value is encoded and read out for these two types of behaviours.

We then tested whether a Bayesian probabilistic representation of value also underlies confidence, assuming conscious access to either the full posterior or a summary statistic of its dispersion. To do so, we tested whether confidence in the value of items would correlate with variability in value ratings. We conducted a second behavioural experiment (Experiment 2) in which 28 participants provided subjective value ratings about the same set of food items as in Experiment 1. However, in Experiment 2, participants were also required to give a subjective confidence rating following each value rating, indicating how confident they were in their estimation of the item’s value (Methods, and Fig. 1a). As in Experiment 1, participants performed two consecutive rating phases of the same set of items, which were used to calculate the individual variability in value ratings for each item (Fig. 1g) as well as the average confidence in value ratings for each item (Fig. 1h). We first replicated the results of Experiment 1, with value ratings being less variable on the edges of the rating scale (𝛽_MixedEffects_ = -0.41 ± 0.03, p < 0.001; Fig. 1g) and average subjective confidence ratings being higher for values closer to the extremes of the rating scale (𝛽_MixedEffects_ = 0.53 ± 0.003, p < 0.001; Fig. 1h). This effect may potentially reflect the compression of probability distributions for extreme values, resulting in lower uncertainty when value space is mapped onto a bounded rating scale^13,48,49^. Crucially, however, subjective confidence was related to the variability in value ratings. Specifically, as expected by our Bayesian inference account, items receiving more variable value ratings were also associated with lower confidence ratings (𝛽_MixedEffects_ = – 0.18 ± 0.06, p = 0.006; Fig. 1i). Similarly, at the participant level, we found that individuals with higher average variability of value ratings were also the subjects that expressed a lower level of confidence in these ratings (𝛽_Robust_ = –3.60 ± 1.2, p = 0.005; Fig. 1j). Importantly, the distance between the mean value assigned by each participant to the set of items and the centre of the scale was also included in the regression, ensuring that the relationship between participants’ variability in value ratings and level of subjective confidence was not an artefact of their distribution of ratings on the rating scale. Hence, these findings extend the results observed with choice consistency to confidence, demonstrating that variability in the estimation of an item’s value is also linked to the subjective confidence in this estimation, both across items and across individuals. Overall, these results replicate and extend previous findings^13^ that have linked preference variability, choice consistency and confidence–an association consistent with predictions from an underlying probabilistic inference of value. This raises the question of how such computations are implemented at the neural level. In the next sections, we directly test whether value representations rely on a probabilistic population code, by analysing fMRI data from the value rating task. Using population receptive field (pRF) encoding models combined with Bayesian decoding, we assess whether this coding scheme is evident in neural activity, and we examine whether the behavioural signatures of probabilistic inference process align with the neural signatures of the probabilistic population code.

### Neural signatures of a linear rate code for value

Using fMRI, we recorded the brain activity of participants during both Experiment 1 and 2 while they were engaged in the value and confidence rating tasks. To confirm that also in our data, mean population activity correlates linearly with subjective value, as has been shown in many earlier studies, we first took a standard General Linear Model-approach and linearly regressed participants’ BOLD signal, at the time they observed each food item, on their corresponding value ratings. This confirmed that activity in the ventromedial prefrontal cortex (vmPFC) correlated positively with subjective value ratings in both Experiment 1 (Fig. 2a) and Experiment 2 (Fig. 2b), in line with previous work^1,4,7,8,50^. We also extended this analysis to the periods following the presentation of food items and found that vmPFC BOLD signal also correlated linearly with value during the subsequent value rating and confidence rating phases (Fig. 2a and 2b). These results replicate classic findings of univariate analyses that have supported a rate code for value and that have suggested a common value signal that may shape behavioural responses at different time points in the experiment. Next, we tested whether brain activity also reflected trial-by-trial confidence judgments about items’ values. We found no significant correlate during stimulus presentation and during the first-order rating period, but a negative relationship between activity in the anterior insula bilaterally and confidence expressed during the second-order rating period (Fig. 2b), again consistent with previous findings^45^. Univariate analyses therefore suggest that a linear rate code in the vmPFC does not capture confidence to the same extent as value judgments, and thus cannot fully explain all behavioural signatures of value inference.

**Fig. 2.**
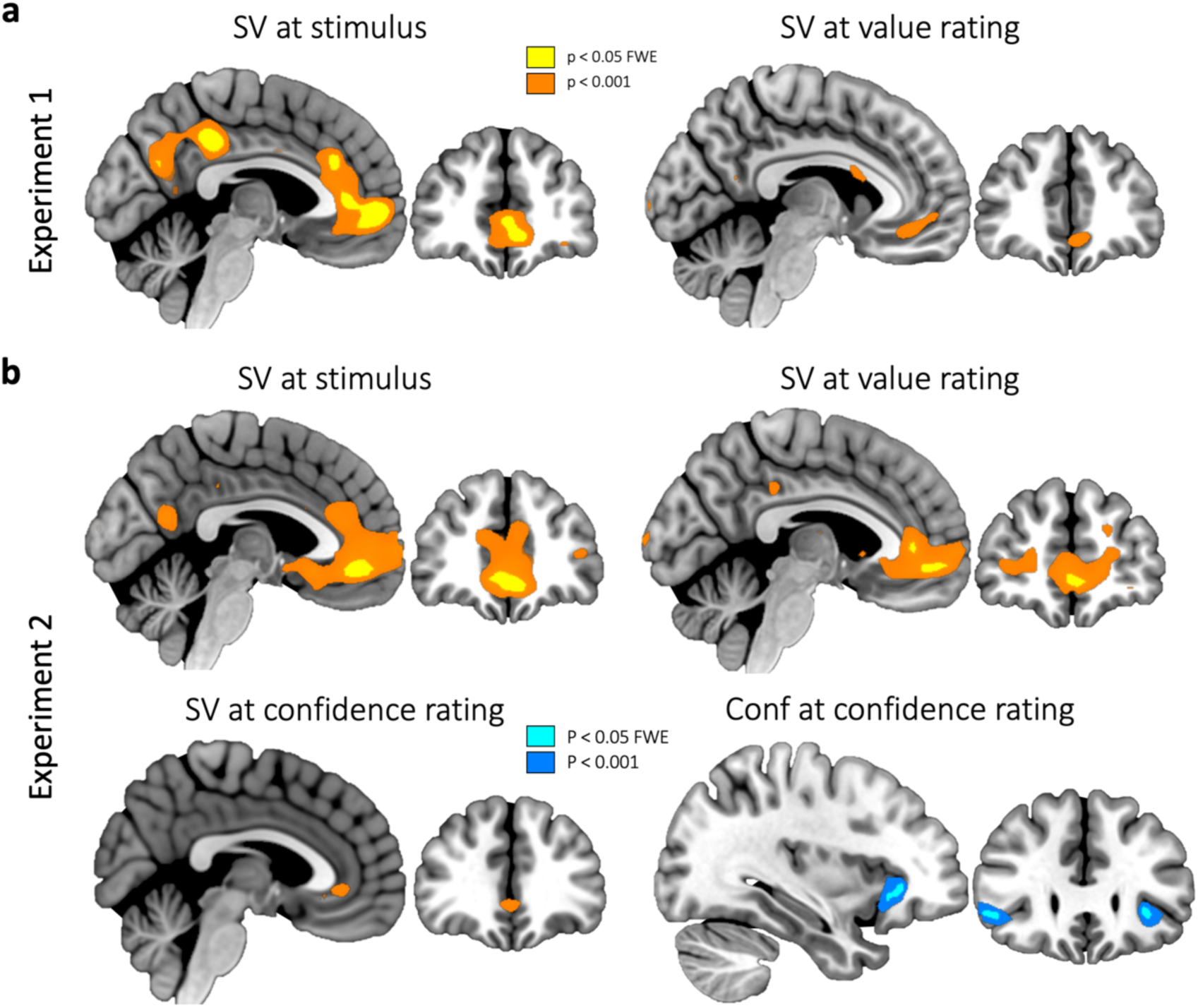
Neural correlates of subjective value and confidence. a,b, Statistical parametric map (SPM) of subjective value (SV) at the time of stimulus presentation and at the time of value rating, in experiment 1 (**a**, n = 36) and experiment 2 (**b**, n = 28). For experiment 2, that comprised a second-order rating, the SPM of SV and confidence (Conf) are also shown at the time of the confidence rating. Colours indicate activations surviving a threshold of P < 0.05, FWE-corrected for multiple comparisons, and P < 0.001 uncorrected, for illustrative purposes.

### Neural signatures of a probabilistic population code for value

To test the novel proposal that OFC/vmPFC uses a non-linear, probabilistic population code for value - similar to that used by sensory cortices for perception^30–33,37,38^ - we first modelled OFC/vmPFC population activity using this coding scheme and decoded both value and its uncertainty as neural signatures of the underlying code. While our primary goal was to decode participants’ subjective value and predict their ratings to test our coding hypothesis, this finding would have broader conceptual implications. Specifically, if subjective value can be experimentally decoded from OFC/vmPFC activity based on a model of the probabilistic population code, it suggests the brain may use similar computational principles to infer value, providing a neural basis for probabilistic models of value inference^13–16^.

In our approach, we assumed that each neural population in the OFC/vmPFC is tuned to a specific magnitude of value and exhibits a Gaussian response profile along the value space (Fig. 3a, top). We fitted such a value population receptive field (pRF) model to participants’ BOLD activity during stimulus presentation, assuming a mixture of neural populations within each voxel. For each participant, we selected voxels best fit by this model within a predefined region of interest (ROI, see Methods and Supplementary Fig. 1) encompassing medial OFC and adjacent vmPFC regions that have both been implicated in value encoding in humans^1–8^. We then inverted the model using a Bayesian decoder, thus simulating decoding by an observer or downstream region, to extract trial-wise indices of value magnitude and uncertainty corresponding to the mean and width of the decoded posterior distribution (Fig. 3a, bottom, see Methods). Crucially, encoding and decoding followed a leave-one-out cross-validation procedure, ensuring robust generalization by fitting the model on all but one run and testing on the held-out run.

**Fig. 3.**
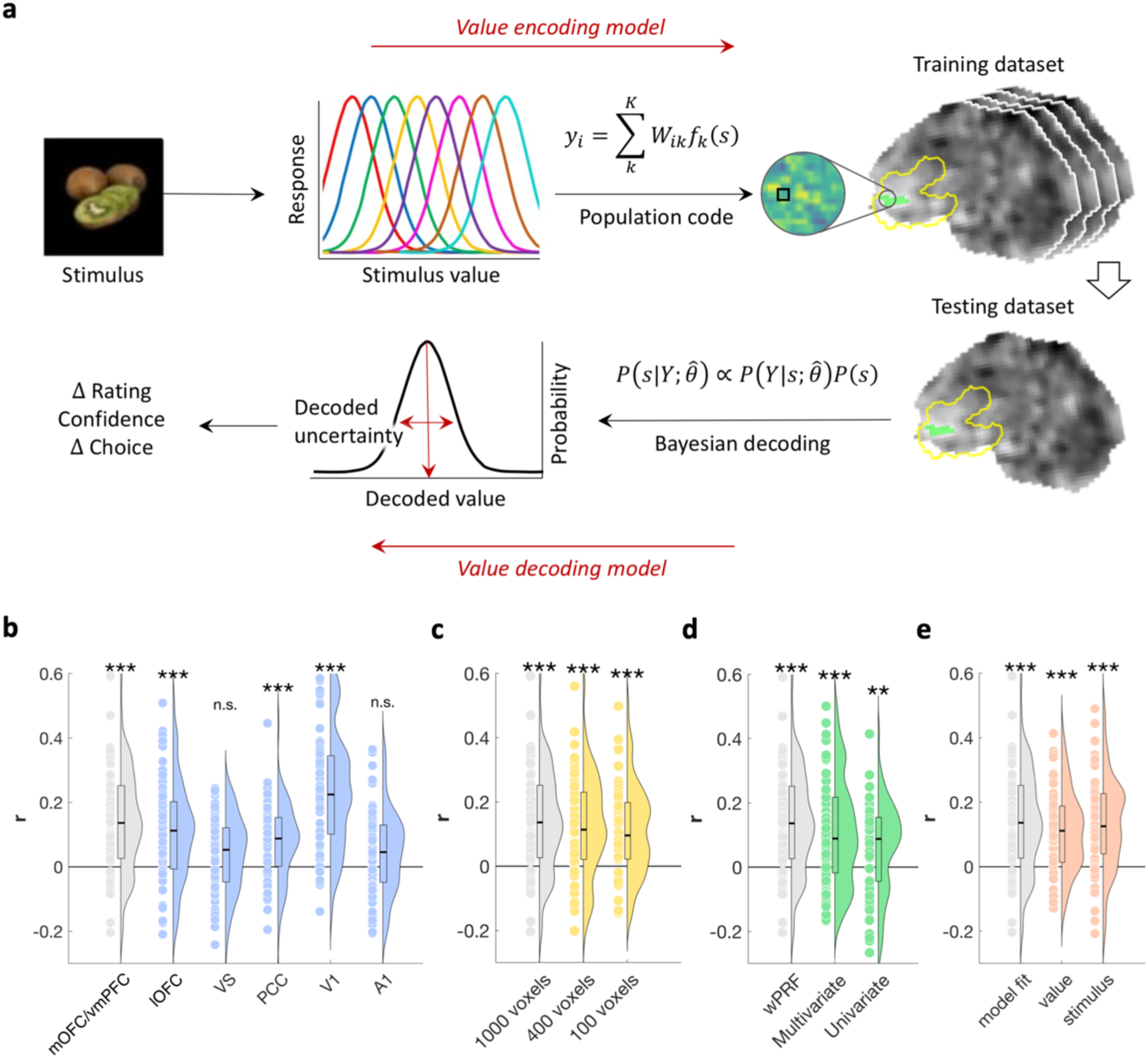
Decoding neural value representation in the ventromedial prefrontal cortex. **a**, Illustration of the value encoding and value decoding models. Observers infer the value of food items, which is encoded in the joint activity of populations of neurons, each tuned to a specific value magnitude. In every voxel 𝑖, BOLD signal 𝑦_𝑖_ is assumed to reflect a weighted sum of responses from 𝐾 neural populations (K = 11) plus neural noise. Population weights B_𝑖𝑘_ that best predicted the BOLD response are estimated in each voxel and each participant, together with noise parameters, in a leave-one-run-out cross-validation procedure, using a training dataset consisting of all but one fMRI run. Value decoding is then performed trialwise on the testing dataset, consisting of the remaining run. Bayesian model inversion of the encoding model yields a posterior distribution over all possible values, given the BOLD activity pattern in the medial OFC/ventromedial PFC. The standard deviation of this posterior is taken as a decoded measure of uncertainty in neural value representations. The expected value of the posterior represents the most likely stimulus value. **b**,**c**,**d**,**e**, Decoding performance. Correlation between observed and decoded value from the leave-one-out cross validation. Grey bars show the averaged correlation across participants using the 1000 voxels in the medial OFC/ventromedial PFC with the best encoding model fit using the weighted pRF model. Coloured bars show the corresponding decoding performance for other regions of interest (blue), number of voxels (yellow), encoding models (green), or voxel selection criteria (pink). Performance for the ventral striatum is only reported for 400 voxels due to the small size of this structure. Error bars represent the s.e.m. across participants. Each dot is a participant. * 𝑃 < 0.05, ** 𝑃 < 0.01, *** 𝑃 < 0.001, after correction for multiple comparisons.

Using this value pRF encoding model combined with a Bayesian decoder, we found that subjective values could be decoded better than chance in the mOFC/vmPFC (subject-level predicted– observed correlations: mean *r* = 0.14, t(63) = 7.1, *p* < 0.001; Fig. 3b). For comparison, the correlation between decoded value and subjective value, as rated by participants, was lower than the correlations reported in previous studies between decoded and observed variables in the perceptual^38^ and numerical domains^34^, but comparable to that observed in neural decoding of working memory^36^. This may reflect lower signal-to-noise ratio and/or greater processing complexity in mOFC/vmPFC value signals than in stimulus-induced responses in sensory and associative cortices. Our Bayesian decoder also tended to outperform both a classical linear multivariate (t(63) = 1.9, *p* = 0.06; Fig. 3d) and a linear univariate approach (t(63) = 3.9, *p* < 0.001) in predicting value ratings.

Decoding of subjective value ratings in the mOFC/vmPFC remained robust across varying numbers of selected voxels, with larger voxel sets yielding increasingly significant results (Fig. 3c). The decoding was also robust when performed on voxels best fitted by the encoding pRF model or on voxels with a highest univariate response to value or to stimulus presentation (Fig. 3e; see Methods). Finally, we tested our value decoder in ROIs of other brain areas implicated in value processing ^7,8^ and in sensory areas such as primary visual and auditory cortices. Value ratings were also decoded above chance level in the lateral OFC (mean *r* = 0.11, t(63) = 5.7, *p* < 0.001; Fig. 3b) and posterior cingulate cortex (mean *r* = 0.07, t(63) = 4.6, *p* < 0.001), although the decoding performance was numerically lower than in the mOFC/vmPFC. Decoding performance was at chance level in the ventral striatum (mean *r* = 0.04, t(63) = 2.9, *p* > 4e-3 for multiple comparisons correction). This may indicate a different value code or a different topographic organization of neuron populations, but may also reflect the smaller size of this brain structure, which may require higher imaging resolution to yield a sufficient number of informative voxels.

To control for specificity, we ensured that subjective value could not be inferred from cortical regions not implicated in the task, such as primary auditory cortex (mean *r* = 0.05, t(63) = 3.0, *p* > 4e-3 for multiple comparisons correction). By contrast, the primary visual cortex V1 showed the highest decoding performance across all ROIs (mean *r* = 0.24, t(63) = 10.6, *p* < 0.001). This result, however, does not necessarily indicate behaviourally relevant subjective value representations in V1. Instead, it may reflect top-down attentional effects associated with increased attention to higher-value items ^51,52^, or responses of V1 populations tuned to visual features associated with value-relevant behavioural goals ^53,54^. Moreover, the greater decoding performance in V1 may result from a higher signal-to-noise ratio in V1 compared to the OFC/vmPFC due to BOLD signal dropout^55^. Irrespective of these considerations, our findings are consistent with our hypothesis in demonstrating that value can be decoded from neural activity, particularly in the mOFC/vmPFC, using a probabilistic population code and a Bayesian decoder.

### Neural signatures of the population code relate to variability in value ratings

Crucially, our probabilistic population model enabled us to decode not only the subjective value magnitude but also its associated uncertainty from neural activity. Because the population code represents a full posterior distribution over subjective value, the precision (or inversely, uncertainty) of this representation can be quantified for each individual trial or item. Specifically, we measured uncertainty as the average standard deviation of the decoded posterior distribution across repeated presentations of the same food item, directly reflecting the variability inherent in the neural population’s probabilistic representation. We first examined whether noisier neural value representations might lead to more variable value ratings for the same option. To do so, we regressed participants’ absolute value rating difference across repetitions of items, both in Experiment 1 and 2, on the decoded neural imprecision, while taking into account the distortion of ratings on a bounded scale by including in the regression the absolute distance between the mean rating for each item and the nearest extreme value of the rating scale. On top of a distortion effect (𝛽_MixedEffects_ = 0.32 ± 0.03, p < 0.001), we found an interaction with decoded neural imprecision (𝛽_MixedEffects_ = 0.066 ± 0.03, p = 0.016, Fig. 4a), indicating that items with more imprecise neural representation of value received more variable ratings across repetitions, with this effect being more pronounced for medium compared to extreme values due to the low variability of ratings on the edge of the scale.

**Fig. 4.**
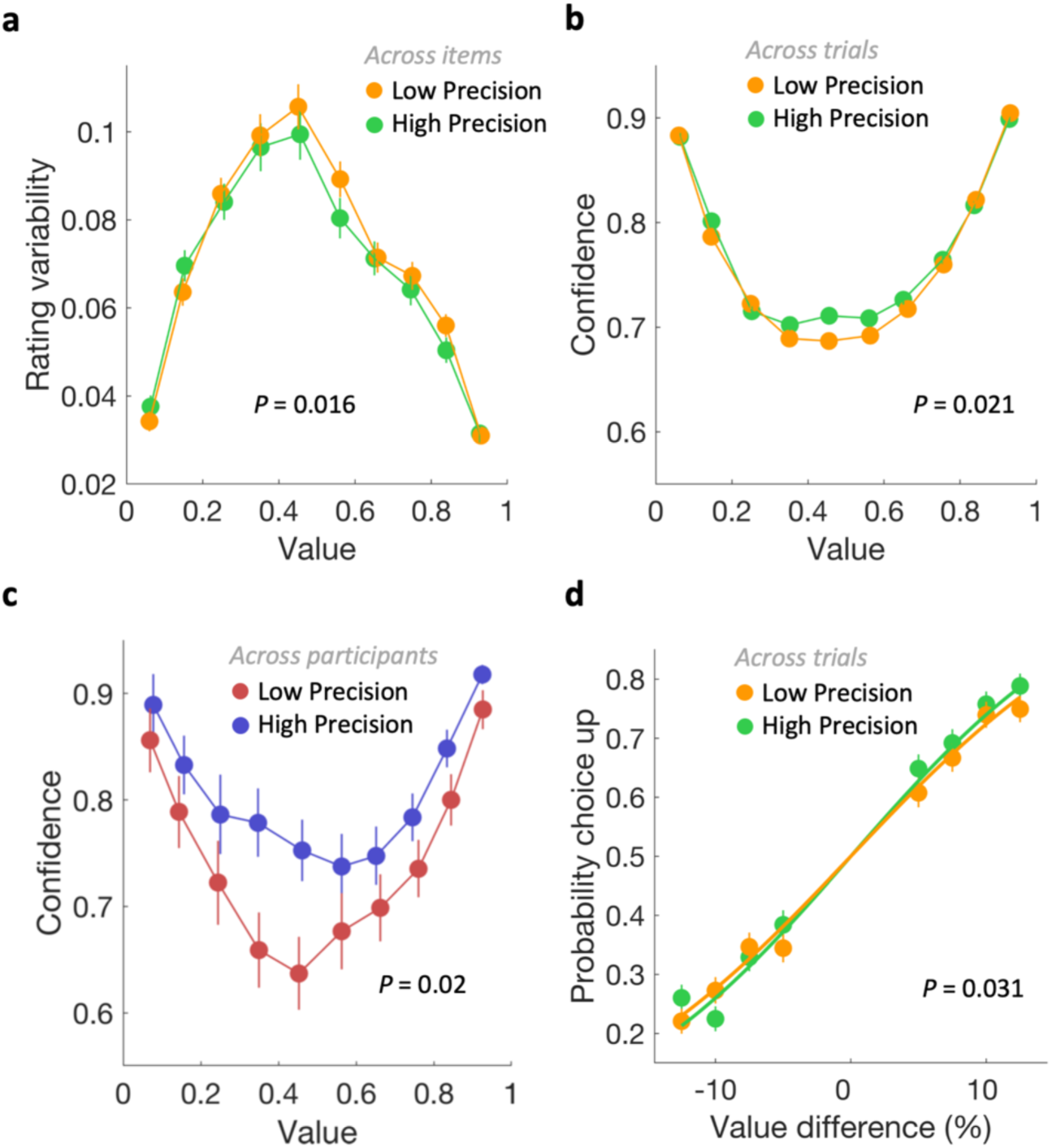
Precision of neural value representation relates to behavioural variability. a, value rating variability plotted as a function of each item’s mean rating (10 value bins) across both rounds in experiment 1. Dots and error bars represent the mean and s.e.m. across items with low (orange) or high (green) decoded neural precision. Items with more imprecise neural value representation exhibit greater rating variability across repetitions. **b,c,** confidence ratings plotted as a function of each item’s mean rating (10 value bins) averaged across both rounds in experiment 2. Dots and error bars represent the mean and s.e.m. across trials (**b**) with low (orange) or high (green) decoded neural precision, and across participants (**c**) with low (red) or high (blue) averaged decoded neural precision. Greater neural imprecision is associated with lower confidence, both across trials and across participants. **d,** Observed choice probability plotted as a function of the two item’s value difference (𝑉_1_ − 𝑉_2_, 8 bins) in experiment 1. Dots and error bars represent the mean and s.e.m. across trials (**d**) with low (orange) or high (green) decoded neural precision. Participants were less likely to choose their preferred item when the alternative was close in value, an effect that was stronger when both neural value representations were imprecise.

These results support the interpretation that rating variability directly reflects the precision of an underlying probabilistic neural value signal. An alternative account is that value is encoded as a point estimate - reflected in the average mOFC/vmPFC activity - and that rating variability arises from moment-to-moment fluctuations in this estimate. To test this, we repeated the regression analysis using the absolute difference in mean BOLD signal between two presentations of the same item (within the same mOFC/vmPFC voxels used for pRF decoding) as a proxy for variability in point-estimates. This analysis revealed no significant association between BOLD fluctuations and rating variability in either Experiment 1 and 2 (𝛽_MixedEffects_, all p > 0.05 for both main effects and interactions). Together, these findings suggest that spontaneous variability in subjective value reports reflect the precision of a probabilistic population code, rather than random noise in point estimate representations.

### Neural signatures of the population code relate to confidence about value

We next tested whether confidence in value judgments might reflect a read-out of the precision of the underlying value representations, by relating confidence ratings to decoded neural imprecision as a key signature of the probabilistic population code. We examined this relationship in Experiment 2 in which participants rated the value of food items and then reported their confidence in those rating. To avoid contamination from response-related signals^48^, we focused on decoded neural imprecision during stimulus presentation only, excluding both rating periods. We regressed the trial-by-trial confidence ratings on decoded neural imprecision, while controlling for distortions in confidence ratings associated with extreme values on a bounded value scale. Specifically, the regression included the absolute distance between of each item’s mean value rating and the nearest scale boundary as a covariate. In line with our hypothesis, we found a significant main effect of distance to the rating scale edge (𝛽_MixedEffects_ = – 0.400 ± 0.02, p < 0.001) and a significant interaction with decoded neural imprecision (𝛽_MixedEffects_ = – 0.03 ± 0.01, p = 0.021). This suggests that confidence judgments may indeed reflect a readout of neural imprecision, since greater neural imprecision was associated with lower confidence, particularly when confidence ratings were not biased by scale boundaries (Fig. 4c).

We then tested whether this trial-by-trial relationship between decoded neural imprecision and confidence also extended across participants. Note that such across-subject analyses were only feasible for the independent confidence ratings (and not variability and choice consistency), as mean decoded neural imprecision is influenced by the variability in value ratings used to train the model (i.e., some of the variance of the decoded posterior may simply reflect observed behavioural, rather than neural variability). Variability and choice consistency—both derived from the same value ratings— could thus not be reliably tested against decoded neural imprecision at the between-subject level. For confidence ratings, we found a significant interaction between distortion due to the bounded scale and participants’ mean decoded neural imprecision (𝛽_MixedEffects_ = – 0.06 ± 0.03, p = 0.02; Fig. 4d), while controlling for mean rating variability. Finally, the relationship between decoded neural imprecision and confidence was specific to core regions of the brain valuation system—the mOFC/vmPFC, lateral OFC, and PCC— and absent in control sensory areas (Supplementary Fig. 2-6). The link between neural precision and confidence complements our earlier findings relating both to variability in value judgments. Together, these results support the view that both preference variability and confidence originate from a readout of the precision of a probabilistic population code in the mOFC/vmPFC

### Neural signatures of the population code relate to choice consistency

Value signals in the mOFC/vmPFC that support value judgments have also been implicated in value-based choices^1,3–8,50^. If the neural signals encoding value with a probabilistic population code in the OFC also guides decision-making, then greater imprecision in this neural representation should also correspond to less consistent choices. This is because the uncertainty encoded in the posterior distribution should reflect the noisiness of the population signal driving the decision. To test this hypothesis, we assessed whether decoded neural imprecision from the fMRI rating task in Experiment 1 predicted participants’ subsequent consistency in choosing the preferred option between two foods outside the scanner. To do so, we regressed participants’ choices on both the subjective value difference between the options and the sum of the decoded neural imprecision of the choice pair. Beyond the expected effect of value difference (𝛽_MixedEffects_ = 0.08 ± 0.01, p < 0.001), we found a significant interaction with total decoded neural imprecision (𝛽_MixedEffects_ = –0.01 ± 0.006, p = 0.031, Fig4e). In other words, across trials, participants were less likely to choose their preferred item when the alternative was close in value, and this effect was stronger when both neural value representations were imprecise (Fig. 4d). This effect was specific to neural imprecision decoded from the mOFC/vmPFC and absent in other tested regions (Supplementary Fig. 2-6). Thus, these findings extend the link between decoded neural imprecision, preference variability, and confidence expressed within the same task, to choices made in a separate session outside the scanner. This does not necessarily imply, however, that the within- and between-task effects reflect the same source of imprecision in neural value representations: Within-task effects may capture trial-by-trial fluctuations in the precision of value representations and could therefore potentially be influenced by attention. In contrast, between-tasks effects may reflect a more stable, item-specific component of value uncertainty, tied to the inherent properties of each food item and the inference process they elicit.

In any case, our results support all key predictions of the probabilistic population code account – namely that the imprecision decoded from mOFC/vmPFC activity relate not only to preference variability and confidence but also to subsequent choice consistency.

### Neural encoding of value in OFC neurons in monkeys

To examine the generality of our findings across species and tasks, we sought to gather empirical support for the value population coding scheme also at the single-neuron level, by examining whether monkey OFC neurons exhibit receptive fields for value. We reanalysed published single-cell data^56^ from two adult male rhesus macaques during an economic choice task. In that experiment, monkeys were presented on every trial with two different choice targets - each associated with the delivery of a distinct fixed juice reward (varying from one to five drops) - and selected from the two their preferred target by pressing the corresponding lever. OFC neurons (n = 1450) were recorded using multi-channel linear recording arrays (Fig. 5a, see Methods, and ^56^ for details). We focused on the 200-400ms period after fixation of the first reward target, when its value was found to be strongly represented in single-cell OFC activity (see shaded area in Fig. 2d in ^56^). We averaged spike counts within this epoch for each trial and fitted different competing models that linked the trial-by-trial response of each neuron to the different value levels using null, intercept, linear, sigmoid, and Gaussian tuning models (Fig. 5b, the latter corresponds to the probabilistic population coding scheme we tested in the fMRI data, see methods for details). For each cell, activity was recorded over 50 to 300 repetitions at each of the five value levels (Supplementary Fig. 7), ensuring robust model estimation. We classified each cell according to the tuning model that had the best log-evidence, but only when this winning model was more likely than the ‘null’ hypothesis of random spikes with a Bayes Factor > 2 (corresponding to “decisive” evidence for this model^57^). 1427 out of 1450 cells could be classified using this method (Fig. 5b). We found that 37% of these cells (n = 529) responded to reward with a constant firing rate across value levels. Consistent with previous reports^19,20,27–29^, about half of the OFC cells (55%, n = 780) showed monotonic responses to value, following either a linear (n = 137) or sigmoid (n = 643) pattern. Approximately half of these monotonic value-coding cells had a positive response to value (n = 414, Fig. 5c and 5d) and half had a negative response (n = 366). Critically, 8% of OFC neurons (n = 118), corresponding to 13% of OFC value-coding neurons, exhibited Gaussian-like receptive fields tuned to a specific value magnitude (Fig. 5e and 5f). The distribution of these cells was approximately uniform across the different reward levels. Note, however, that the proportion of neurons with Gaussian-like tuning may have been somewhat underestimated, as the restricted range of reward values in that experiment may in principle have led to a misclassification of such neurons as monotonic. In any case, these findings demonstrate that monkey OFC contains a subpopulation of neurons with non-linear value tuning — including bell-shaped and sigmoidal response profiles — consistent with the kind of heterogeneous tuning that supports probabilistic population codes. This provides single-cell evidence broadly compatible with the probabilistic inference framework underlying our fMRI decoding model, which implements a specific instantiation of this principle using Gaussian tuning functions.

**Fig. 5.**
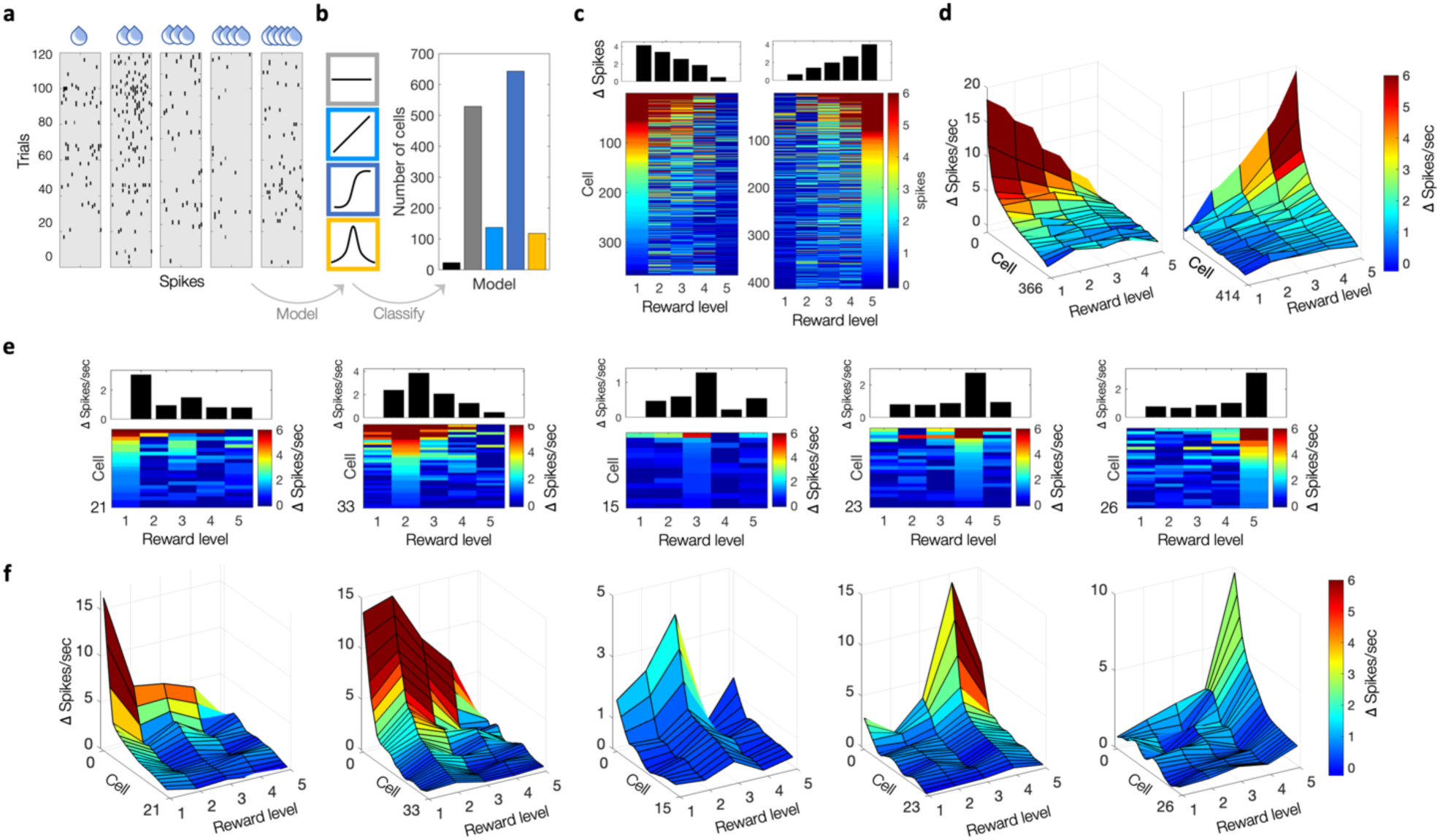
Neuronal encoding of value in the monkey orbitofrontal cortex. Data are taken from McGinty & Lupkin, 2023. Single-cells recordings of neurons (n = 1450) in the orbitofrontal cortex obtained from two macaque monkeys during an economic choice task. Neural activity spans 200-400ms after the monkey fixated on the first of two reward cues, each associated with five possible levels of juice reward (see Fig. 2d in McGinty & Lupkin, 2023). **a,** Raster plots of an example cell (cell N° 1323) across five value levels. **b,** Cell classification using Bayesian model comparison across five response models: null (black), intercept (grey), linear (light blue), sigmoid (dark blue), Gaussian-like (yellow). Non-classified cells (Bayes factor <2 for the best model compared to the null model) are shown in black. **c-d,** Neuronal activity patterns for cells showing positive and negative linear or sigmoid responses. **c,** Top: Neuronal activity averaged across cells and reward level. Bottom: Neuronal activity shown individually per cell. Cells are sorted by their peak firing rate. **d,** Similar to **c** (bottom), but smoothed using a moving average of 20 cells. **e-f,** Neuronal activity patterns for cells showing Gaussian-like responses. Cells are grouped according to the value level to which they respond most strongly. **e,** Top: Neuronal activity averaged across cells and reward level. Bottom: Neuronal activity shown individually per cell. Cells are sorted by their peak firing rate. **f,** Similar to **e** (bottom), but smoothed using a moving average of 5 cells.

## DISCUSSION

Value-based decision-making is traditionally thought to rely on value signals encoded as point estimates via a linear rate code. In this framework, variability is typically attributed to neural noise, either disrupting response selection^58^ or corrupting value signals themselves^59–61^. Yet, fully random neural noise should average out across the population of neurons and cannot explain various aspects of behavioural variability in decision-making. Here, we offer the alternative perspective that value is encoded through a probabilistic population code. This scheme allows neural activity to convey both the estimated value and its associated uncertainty to connected regions, providing a principled account of how the brain may estimate both value and the associated uncertainty for decision-making. We provide empirical evidence that both key signatures of a probabilistic population code - decoded value and uncertainty - are reflected in neural data. First, using a model implementing such a coding scheme with Bayesian decoding, we could decode subjective value from the mOFC/vmPFC above chance level. Second, a core prediction of Bayesian models of value inference — for which a probabilistic population code could provide a neural substrate — is that preference variability, choice stochasticity, and confidence may stem from a shared probabilistic uncertainty, resulting in correlated fluctuations across items. Consistent with this, we found that all these behavioural signatures of probabilistic value inference were not only correlated with each other, but also with the uncertainty in value readouts from fMRI activity. This supports the view that similar population coding principles may govern lower-level perceptual and cognitive representations in sensory and association cortices^30–40^ and higher-level subjective decision variables in value-related brain systems^13,62^. Our findings thus offer a unified perspective on value-based preference, choice variability, and confidence, suggesting they should be studied not in isolation but as interconnected outcomes of a common underlying neural coding scheme. Our results challenge the traditional view that value is represented exclusively as a scalar quantity (i.e., a point estimate), via a linear rate code based on the averaged activity of neural populations. Instead, they support an additional probabilistic code scheme in which populations of neurons tuned to a different value magnitude jointly encode value, allowing both value and uncertainty to be decoded from the population response ^30,31,40,62,63^. While noisy point estimates of value could, in principle, be the readout of value inference processes characterized here and in prior work^13^, our findings provide a critical distinction. Specifically, we show that uncertainty decoded directly from single neural value representations – rather than inferred from their variability across repetitions – predicts imprecisions in preferences, choice consistency, and confidence. This challenges the prevailing idea that noise in point estimates is the primary source of variability in value-based decision-making and instead supports a probabilistic population coding scheme as a more comprehensive account.

The ability to decode value uncertainty from neural activity raises the questions about its origin. Our finding that uncertainty decoded in the value rating task performed in the scanner predicts choice stochasticity in a separate task outside the scanner suggests that some of this uncertainty reflects a stable, item-specific property. Multiple components of the value inference process may contribute to it. According to a prominent theory, the brain constructs subjective value by integrating an option’s features^9,10^. However, the precision with which these features are represented likely varies across individuals. For instance, recent work has shown that risk aversion for monetary options is associated with the acuity of numerical representations in the parietal cortex^34,46^. Additionally, memory-guided decisions may be impaired by imperfect recall of past experience^64^, leading to inaccurate retrieval of item features and thus imprecise value estimation. Finally, limited prior exposure to an item can introduce irreducible uncertainty in linking features to value^65^.

However, our findings do not imply that variability in decision-making stems solely from probabilistic uncertainty tied to each option’s specific features. Rather, the precision of neural value representations may also be shaped by broader cognitive processes engaged during value inference. Prior research has highlighted the roles of attention^66,67^, working memory and cognitive control^68,69^ in shaping value-based decision-making. These factors can drive moment-to-moment fluctuations in neural precision, as seen in the context of working memory^36^. By accumulating more evidence about reward, decision makers may average out such fluctuations, consistent with findings that repeated evaluation of the same item reduces uncertainty, lowers rating variability, and increase choice consistency^70^. Additional sources of imprecision have been identified, including exploration strategies^61,71,72^, fluctuations in OFC/vmPFC activity^59–61^, and response noise^58^. Still, the precision of neural value representations may represent a fundamental coding property that integrates both item-specific uncertainty and broader inference-related noise linked to limited cognitive resources. In any case, the probabilistic framework we propose here provides a unified account of how both internal and external factors may contribute to variability in value-based decision-making.

Our finding that the precision of neural value signals correlates with subjective confidence support the idea that humans have conscious access to (at least a summary statistic of) the population activity reflecting uncertainty about value. This aligns with evidence from perception, where confidence in in estimating visual features (e.g., stimulus orientation) has been linked to the imprecision of decoded sensory information represented as a probability distribution^45,73^. Our results are also consistent with Bayesian theories^42–44^ proposing that confidence in making the right choice is computed from the posterior probability of being correct given the available evidence. Whereas most prior research on confidence in value-based decision-making has focused on two-option choices^74,75^, our study examined confidence in estimating the value of a single option^13,48^. Nonetheless, the probabilistic coding framework used here could naturally extend to decision contexts. In a choice scenario, the relevant decision variable — and basis for computing confidence — would be the value difference between options, a quantity previously linked to mOFC/vmPFC activity^76–78^. Finally, while our results suggest that uncertainty is reflected in mOFC/vmPFC activity, it remains unclear whether this information is consciously available directly from population activity. More plausibly, the brain may rely on downstream regions such as the dorsal anterior cingulate cortex, the anterior insula, or the rostrolateral prefrontal cortex^45^, to extract and interpret the imprecision in value representations to support computations like confidence.

Finally, we provide empirical support for a probabilistic population coding scheme at the single-neuron level in monkey OFC. In line with previous reports^19,20,27–29^, most value-coding neurons exhibited monotonic response profiles – at least within the range of values that were presented to them in the experiment. Similar monotonic coding has been observed in sensory and sensorimotor regions^79–84^. Crucially, we identified a subset of OFC neurons with non-linear, bell-shaped (i.e., Gaussian-like) tuning curves, a coding scheme extensively reported in perceptual and motor cortices^85–88^ as well as in hippocampal place cells^89^ and previously observed in the monkey OFC and ACC in similar proportions to those reported here^90^. Our results further suggest that these two coding schemes, monotonic and Gaussian-like, coexist in the OFC, as previously observed in occipital and parietal cortices^91–93^. This raises the possibility that these coding schemes may serve distinct functional roles. While sigmoid and Gaussian tuning are computationally related^84^ (e.g., sigmoid functions can be combined to reconstruct tuned responses) and can both support probabilistic population codes, they differ in their functional implications: Gaussian tuning may support minimal decoding time^94^ and local generalization of learning^95^, whereas monotonic responses may promote global generalization^96^. Moreover, these coding schemes may reflect adaptations to different behavioural demands, facilitating either rapid flexible responses or vigorous actions, respectively^97^. Overall, the coexistence of tuning types in the OFC may reflect behavioural constraints and serve to optimize coding efficiency. More broadly, the heterogeneity of neuronal response profiles has been shown to enhance population coding accuracy^98^, with even untuned neurons contributing to information encoding under certain conditions^99^. While the presence of tuned value-coding neurons in OFC remains to be further confirmed, especially in humans, our findings offer important implications for models of decision-making in neuroscience, psychology, and neuropsychiatry.

In conclusion, our results support a probabilistic population code for value in the human mOFC/vmPFC, implemented through non-linear response profiles (Gaussian tuning in our model), and provide preliminary single-neuron evidence for similar coding principles in monkey OFC. This probabilistic population code allows the uncertainty associated with subjective value to be directly decoded from mOFC/vmPFC population activity. By representing a full posterior distribution over value, this code mechanistically explains key behavioural signatures of imprecision in value-based decisions, such as preference variability and stochastic choice patterns. Importantly, neural imprecision also correlates with confidence, suggesting that humans can access and report the imprecision in their own value representations. Together, these findings provide a plausible neural substrate for recent Bayesian theories of value inference^13–16^ and a unified framework that links preferences, choices, and confidence to the precision of neural value representations. This framework may shed light on how the brain integrates external uncertainty about outcomes in the environment with internal uncertainty arising from limited computational precision and may help explain systematic biases in human decisionmaking.

## METHODS

### fMRI EXPERIMENTS

#### Participants

64 healthy young volunteers participated in this study (age 19–40 years; *n* = 36 in experiment 1, 19 females; *n* = 28 new participants in experiment 2, 11 females). Participants were instructed about all aspects of the experiment and gave informed consent prior to participating. None of the participants suffered from any neurological or psychological disorder or took drugs or medication that interfered with participation in our study. We also screened participants for MR compatibility before their participation in the study. The experiments conformed to the Declaration of Helsinki and the experimental protocol was approved by the Ethics Committee of the Canton of Zurich.

#### Procedure

All fMRI experiments were conducted at the Laboratory for Social and Neural Systems Research of the Department of Economics, University of Zurich. Participants completed the consent forms and MRI screening on their arrival. Before performing the tasks, participants were given written instructions about the tasks. Participants of experiment 1 read the instructions for the value rating tasks and the choice task. Participants of experiment 2 read the instructions for the value-and-confidence rating task. Participants performed practice trials of both tasks before they were brought to the MRI scanner room. Then, participants of experiment 1 performed the value rating task and participants of experiment 2 performed the value-and-confidence rating tasks inside the MRI scanner, where we recorded their behavioural and neural measures of value-based preferences. Participants of experiment 1 subsequently performed the choice task outside the MRI scanner, in a behavioural testing room. For all experiments, participants were asked not to eat or drink anything for 3 h before the start of the experiment. All experiments took place between 09:00 and 17:00. After the experiment, participants were required to stay in the room with the experimenter while eating the food item that they had chosen in a randomly selected trial of the choice task (see below). Participants also received monetary compensation for their participation in the experiment.

#### Value rating task

Participants of experiment 1 were asked to provide subjective-preference ratings for a set of 64 food items using an on-screen slider scale (Fig. 1a). Participants were informed that all food items were available in our lab. Importantly, they saw all food products before the rating tasks so that they could effectively use the full range of the rating scale. Food items were selected based on previous studies^13^ to cover the full range of subjective values people typically assign to food items, from options most find unappealing (for example, raw broccoli) to those almost everyone finds highly appetitive (for example, ice cream). Items were displayed in the centre of the screen with a duration of 2500 ms. The rating scale appeared only once the image disappeared from the screen, and the participants were instructed to indicate ‘how much they liked the presented food item’ as fast as possible within a 5000 ms time window. The slider scale was continuous with no numbers displayed and the initial location of the slider was randomized for each item to reduce anchoring effects. Participants were informed that the right end of the scale would indicate items that they liked the most, while the left end would indicate items that they disliked the most. The task comprised two ratings of the same set of 64 food items, in two consecutive rating phases. Phase 2 of the rating task mirrored phase 1 and occurred directly following phase 1. The sequence of item presentation was randomized in each phase. Importantly, participants were unaware prior to phase 1 that a subsequent rating phase would follow. This precaution was taken to prevent participants from deliberately memorizing the slider’s position during phase 1, ensuring an unbiased measure of variability in the value estimates.

#### Choice task

Participants of experiment 1 were asked to choose, outside the MRI scanner, between pairs of food items taken from the set of items presented in the rating tasks. An algorithm selected a balanced set of choice trials divided into four value difference levels on the rating scale (rating difference ∼5%, ∼7.5%, ∼10%, and ∼12.5% of the length of the rating scale), as defined by the average rating across phases 1 and 2 provided by each participant. Choice trials started with a fixation cross displayed in the centre of the screen for 1.7–2.5 s. Two food items were then displayed simultaneously, one in the upper and one in the lower part of the screen. (Fig. 1b). The food items were displayed until participants responded and they had up to 4 s to make a choice. Participants were asked to choose which of the two items (upper or lower) they would prefer to eat at the end of the experiment. Participants responded by pressing one of two buttons on a standard keyboard with their right-index finger (upper item) or their right thumb (lower item). We defined a consistent choice as a trial in which the subject chose the item with a higher mean rating in the rating task. The task comprised 176 trials divided into 4 runs of 44 trials each. The trials were fully balanced across rating-difference levels and location of consistent response option (up or down).

#### Confidence rating task

Participants of experiment 2 performed the value-and-confidence rating task. This task was identical to the value rating task, except that participants indicated after each value rating their confidence in their rating (Fig. 1a). The confidence scale appeared after a 5100 ms interval following the value rating. Participants were instructed to indicate their confidence as fast as possible within a 4500 ms time window. We informed participants that the right end of the scale would mean ‘not at all’ confident, while the left end would mean ‘totally’ confident.

#### Behavioural analyses and statistics

Preference-rating variability in experiments 1 (n=36), and 2 (n=28) was computed as the standard deviation for each item across the two rating phases. For illustration purposes, we plotted rating variability as a function of the mean rating (Fig. 1c,g). To investigate the influence of extreme values on rating variability, we performed a hierarchical linear mixed-effects regression of rating variability on the distance between each item’s mean rating and the centre of the rating scale. We employed the same hierarchical linear mixed-effects regression model on confidence in experiment 2, defined as the average confidence rating for each item. Similarly, we performed a hierarchical logistic mixed-effects regression of the consistency of choices on three regressors of interest, namely: value difference, summed-variability (Var, defined as the sum of the sum of the two standard deviations of the two food items presented in each trail), overall value (OV, defined as the sum of mean-rating values of the two food items presented in each trial). All regressors of interest were included in the same model. All mixedeffects models in this study had varying subject-specific intercepts (that is, we performed random-effects analyses) and were conducted using the *fitlme* function implemented in Matlab (Mathworks). Hypothesis testing for parameters of the regression model was performed using two-sided t-tests.

To investigate the influence of the precision of neural value signals on decision-making (Fig. 4), we conducted similar mixed-effects models with decoded neural imprecision (see fMRI decoding model below) as regressor of interest. For preference-rating variability, the model included the level of decoded neural imprecision averaged across the two presentations of each item. For confidence ratings, it included the decoded neural imprecision estimated on each trial. For choice consistency, the model included the sum of the mean decoded imprecision of the two food items presented in each trial. To complement the item-by-item and trial-by-trial analyses, we performed a between-participants analysis (Fig. 4c), regressing confidence ratings against participants’ mean decoded neural imprecision. Hypothesis testing for the regression model parameters was conducted using one-sided t-tests, based on the a priori hypothesis that higher neural uncertainty translates into more imprecise behaviour. While statistical tests assess the continuous effect of the neural value signal precision, Fig. 4 illustrates behavioural effects for high versus low decoded precision (median split by trials in 4a, items in 4b/d, and participants in 4c).

#### MRI acquisition and preprocessing

We acquired functional MRI data using a Philips Achieva 3T whole-body MR scanner equipped with a 32-channel MR head coil. Specifically, we collected eight runs with a T2*-weighted gradient-recalled echo-planar imaging sequence (123 volumes + 5 dummies in Experiment 1; 158 volumes + 5 dummies in Experiment 2; flip angle 90°; repetition time, TR = 2,625 ms; echo time, TE = 30 ms; matrix size 96 × 96, field of view 240 × 240 mm; in-plane resolution of 2.5 mm; 40 slices with thickness of 2.5 mm and a slice gap of 0.5 mm; SENSE acceleration in phase-encoding direction with factor of 1.5; time of acquisition 5:38 min in Experiment 1, 7:12 min in Experiment 2). Additionally, we acquired high-resolution T1-weighted 3D MPRAGE image (field of view 256 × 256 × 181 mm; resolution 1 mm isotropic; inversion time, TI = 2,800 ms; 256 shots, flip angle 8°; TR = 8.3 ms; TE = 3.9 ms; SENSE acceleration in the left to right direction 2; time of acquisition 5.57 min) for image registration during post-processing. Pre-processing was performed with fMRIPrep v.1.5.0^100^ using standard settings. For more information on pre-processing, see the Supplementary Information.

#### Neuroimaging analysis

We first estimated a general linear model (GLM) to generate statistical parametric maps (SPMs) of the subjective value of food items. All trials of the rating tasks were modelled with the following regressors: first, box-car functions were used to model the different epochs within each trial - stimulus display (2500 ms), value rating epoch (5000 ms), and confidence rating epoch (only in experiment 2, 4500 ms). Second, the value and confidence ratings provided by each participant were incorporated as parametric modulators for all three epoch regressors. All regressors were convolved with a canonical hemodynamic response function (HRF) before regressing the BOLD signal in each voxel onto them. Linear contrasts of regression coefficients (betas) were first computed at the individual participant level, in the MNI space, and then taken to a group-level random effect analysis (using one-sample t-test). All reported significant activations contained voxels surviving a threshold of p< 0.05 after familywise error (FWE) correction for multiple comparisons, otherwise mentioned. We also estimated the model in native space for each participant. This was done to allow selecting voxels whose activity best correlated with value, in native space. These voxels were used as input for the value decoding pRF model (see below).

We then estimated a second GLM to estimate neural activity at each trial during value estimation. Each trial was modelled with a distinct box-car function over the value estimation epoch (2500 ms) and convolved with a canonical HRF. The reliability of single-trial response estimates was improved by identifying an optimal HRF at each voxel, optimizing the set of GLM nuisance regressors (GLMdenoise technique), and applying a custom amount of ridge regularization at each voxel, using the GLMsingle toolbox (github.com/cvnlab/GLMsingle). Single-trial response estimates were then used as input to the value encoding and decoding pRF models (see below).

Decoding was conducted on voxels selected within anatomically defined masks (Supplementary Fig. 1), using the MarsAtlas-Colin27-MNI cortical parcellation atlas (meca- brain.org/software/marsatlas-colin27) for cortical structures, and the Harvard-Oxford atlas (fsl.fmrib.ox.ac.uk/fsl/fslwiki/Atlases) for subcortical structures. The following atlas labels were applied: medial OFC/ventromedial prefrontal cortex (bilateral ‘Ventromedial Orbito Frontal Cortex’, ‘Ventromedial Prefrontal Cortex’ and ‘Anterior Cingulate Cortex’), lateral OFC (bilateral ‘Ventrolateral Orbito Frontal Cortex’ and ‘Ventral Orbito Frontal Cortex’), PCC (bilateral ‘Posterior Cingulate Cortex’), primary auditory cortex (bilateral ‘Caudal Superior Temporal Cortex’), primary visual cortex (bilateral ‘Caudal Medial Visual Cortex’), and ventral striatum (bilateral ‘Accumbens’). All masks underwent dilation using the Scipy library’s ‘binary_dilation’ function (scipy.org) with five iterations and were subsequently mapped onto each participant’s native space using the ANTS library’s ‘applyTransform’ function (nitrc.org/projects/ants).

### Value encoding model

To model the BOLD response to stimulus value, we used a pRF generative model assuming that neurons in the brain valuation system are selective to value magnitude. More specifically, the model assumes that the BOLD response 𝑦_𝑖_ of voxel 𝑖 to a stimulus 𝑠 reflects a weighted sum of responses from 𝐾 neural populations (𝐾 = 11), each tuned to a specific value magnitude^38^. The responses of the neural population are modelled by Gaussian tuning curves 𝑓_𝑘_, regularly spaced over the value space in increments of 0.1 within the range [0, 1], with a standard deviation (sd) of 0.075. The contribution of each neural population is determined by a weighting parameter B_𝑖𝑘_:

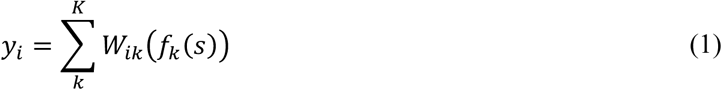

We modelled BOLD data for each voxel and individual, using a gradient descent optimization, implemented in TensorFlow (www.tensorflow.org), to find the population weights B_𝑖𝑘_that best predicted the BOLD response in each voxel. All parameters of the model were estimated jointly by maximizing their likelihood given the stimulus value. The resulting Python package (braincoder) can be found on GitHub (https://github.com/Gilles86/value_prf).

### Value decoding model

To decode neural value representations, we implemented a Bayesian inversion of the value encoding model, building on previous work on encoding-decoding models^34,38^. Crucially, model parameters were estimated using the fMRI BOLD responses to stimulus presentation in a leave-one-run-out cross-validation procedure. The generative model was fitted to a training dataset, consisting of data from all but one fMRI run. The estimated model parameters were then tested on the testing dataset, consisting of the held-out run. The univariate encoding model 𝜙(𝑠) was extended with a multivariate Gaussian noise model, I, to estimate a conditional probability distribution over a multivariate response pattern, 𝑌 = [𝑦_1_, …, 𝑦_𝑛_], given stimulus value, 𝑠:

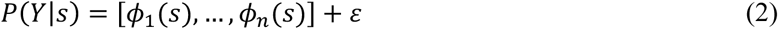

where:

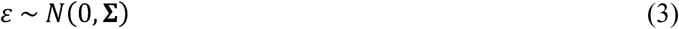

Building on previous work^34,38^, the covariance structure S of the noise model was defined as:

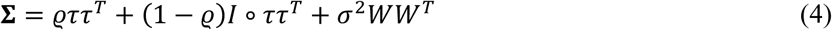

where 𝜚 ∈ [0,1] is a scalar that quantifies noise correlation across voxels; 𝜏 is a vector containing the standard deviation of the residuals in each voxel; 𝐼 is the identity matrix; 𝜎^2^ is a scalar that specifies the variance of noise shared across neural populations of similar value preference; and BB^𝑇^ a matrix that quantifies the amount of overlap in neural populations of different voxels.

We implemented the noise model in Tensorflow and used gradient descent to estimate ρ, 𝜏, 𝜎2 with a maximum likelihood cost function. We fixed the parameters 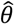 of the encoding model to the values estimated in the training phase. Fitting the noise model enabled us to decode the imprecision of value representations, measured as the width of the posterior probability distribution 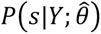, which represent the probability of all possible value magnitudes 𝑠, given the BOLD data 𝑌∗ of a given trial of the held-out run.

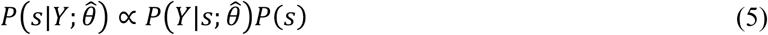

We used the mean of this posterior 𝐸[𝑠] to predict the value 𝑠 of the unseen data 𝑌∗, and the standard deviation of the posterior to quantify the associated uncertainty of value representations on a trial-to-trial basis.

To benchmark the value pRF Bayesian decoder, we implemented two simpler decoding approaches using the same single-trial beta estimates and cross-validation procedure. For the univariate decoder, single-trial beta estimates for selected voxels were averaged into a scalar mean activation per trial, and ordinary least square regression was used to predict subjective value ratings on held-out runs. For the multivariate linear decoder, we replaced the Gaussian pRF encoding model with a linear model (LinearModelWithBaseline, braincoder), where the predicted response in voxel v is:

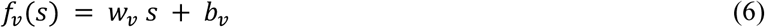

with response weights *w_v_* fitted via L2-regularized ridge regression (α = 0.1) and baseline *b_v_* set to the mean training-run activation. Decoding then followed the same Bayesian inversion procedure as the pRF decoder: the residual covariance structure (ρ, τ, σ²) was estimated on training runs, and the posterior 𝑃(𝑠 | 𝑌 ∗) computed over 150 linearly spaced bins spanning the full value range, with decoded value taken as the posterior mean E[s]. For all three decoders, performance was quantified as the Pearson correlation between predicted and reported value ratings, averaged across folds.

To assess decoder robustness, we compared three voxel selection methods: (i) selecting the N voxels best fit (highest R^2^) by the pRF model using training data only to ensure independence from test data ("model fit”); (ii) selecting voxels with the highest value-related t-statistics from the first GLM with value ratings as a parametric modulator (“value”); and (iii) selecting voxels with the highest value-related t-statistics from the first GLM with stimulus presentation as an event regressor (“stimulus”).

### Single-cells recordings in monkey OFC

Single-cell data are taken from McGinty & Lupkin, 2023. Neural unit signals were obtained from two adult male rhesus macaques K and C, using multi-channel linear recording arrays (Plexon-V-Probes) with 16, 24 or 32 channels, spaced 50 µm or 100 µm apart. OFC was identified on high-resolution MRI scans from each animal. Cells were isolated using semi-automated spike-sorting procedures (see McGinty & Lupkin, 2023 for detailed methods). 848 cells were isolated for monkey K; and 602 cells for monkey C. Neural activity was recorded during an economic choice task. On each trial, monkeys were shown two choice targets, selected from 12 unique targets, each associated with the delivery of a fixed juice reward between one to five drops. After an initial fixation period, monkeys were free to view the two choice targets by self-paced fixations, and to make a choice at any time by lifting and pressing two levers. The target stimuli and display were designed so that the targets were not visible to the monkey until the monkey fixated directly upon them (see McGinty & Lupkin, 2023 for detailed methods). Juice reward was delivered via a gravity-fed reservoir and solenoid valve.

Spike count data were time-locked to the viewing of the first reward target and averaged across trials for each cell within a 200-400ms period after fixation of the first target (corresponding the grey shaded region in Fig. 2d in McGinty & Lupkin, 2023). Each cell was recorded in at least 50 to 300 trials for each of the five reward levels (Supplementary Fig. 7).

Individual cell activity patterns of response to the different value levels were modelled as intercept, linear, sigmoid, or Gaussian tuning functions, and compared to a null model. Under the null model, the spike averaged activity 𝑠 consisted of random noise 𝑠 = I, where I ∼ Γ(𝑘, [) and 𝑘 and [are the shape and the scale parameters of a Gamma distribution, respectively. The intercept model 𝑠 =𝛽 + I, and the linear model 𝑠(𝑣) = 𝛽 + 𝛼𝑣 + I, included an intercept 𝛽 and slope 𝛼 related to the reward value 𝑣. Sigmoid responses were modelled as 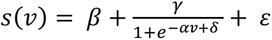, where 𝛼, 𝛽, f, g are the slope, baseline, inflection point, and maximum of the sigmoid, respectively. Gaussian tuning curves corresponded to the model used for the analysis of the fMRI data and were modelled as 𝑠(𝑣) = 𝛽 +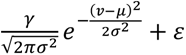, where 𝛽, 𝜇, 𝜎, g are the baseline, mean, standard deviation and maximum of the curve, respectively. The different models were inverted using a variational Bayes approach under the Laplace approximation, implemented in the VBA toolbox (mbb-team.github.io/VBA-toolbox). This algorithm not only inverts nonlinear models with efficient and robust parameter estimation, but also estimates the model evidence, which represents a trade-off between accuracy (goodness of fit) and complexity (degrees of freedom). The following non-informative priors were used for parameter estimation: shape 𝑘 = 1 and scale [= 1 for the noise model; slope 𝛼 = 0 and intercept 𝛽 = 0 for linear functions; baseline 𝛽 = 0, slope 𝛼 = 0, max g = 5, and inflection point f = 2.5 for sigmoid functions; baseline 𝛽 = 0, mean 𝜇 = 2.5, standard deviation 𝜎 = 5, and maximum g = 5 for gaussian functions; s.d. was set to 10 for all priors. The model log-evidence was then used to classify cells based on the model that best accounted for neuronal tuning curves to value in the monkey OFC, provided it was more likely than the ‘null’ hypothesis of random spikes (Bayes Factor > 2). Cells showing linear or sigmoid responses were further grouped as positive or negative responding cells, while cells showing Gaussian-like responses were grouped according to the value level to which they responded most strongly.

For illustration purposes, the raw spike count was averaged across each type of responding cell and for each value level. To account for heterogeneity in cells response ranges, the spike count was baseline corrected, removing the mean activity of the lowest response across value-levels for each cell. Population activity patterns were illustrated by sorting cells of each type by their baseline-corrected peak firing rate and by smoothing firing rate using a moving average of 20 (linear and sigmoid cells) or 5 cells (Gaussian-like cells).

## DATA AVAILABILITY

The behavioural data used in this study are available at https://github.com/ruffgroup/value_prf. The neuroimaging data are available at https://openneuro.org.

## CODE AVAILABILITY

Analysis code is available at https://github.com/ruffgroup/value_prf.

## ACKNOWLEDGEMENTS

We are grateful to C. Schnyder, K. Treiber and M. Moisa at the. Zurich Center for Neuroeconomics for their excellent assistance in recruitment and participant facilitation. This work received funding from the Marie Skłodowska-Curie Actions (MSCA) postdoctoral individual fellowship program (grant no. 890141 to R.L.B), the Bettencourt Schueller Foundation (grant to R.L.B), the Swiss National Science Foundation (SNSF) (SPARK grant no. CRSK-3_190501 to R.L.B, and grants no. 105314_152891 and 10006863 to C.C.R), and the University Research Priority Program ‘Adaptive Brain Circuits in Development and Learning’ at the University of Zurich. G.d.H. was funded by the Dutch Research Council NWO (Rubicon grant no. 019.183SG.017/8O3B) and the University of Zurich (Forschungskredit grant no. K-33153-02-01). The funders had no role in the study design, data collection and analysis, decision to publish or preparation of the manuscript.

## AUTHOR CONTRIBUTION

R.L.B, G.d.H, M.G., R.P and C.C.R. developed the experimental design and procedures and contributed to the analysis pipeline of the behavioural data. M.G. and R.P. collected the behavioural and imaging data. R.L.B. and G.d.H. set up the analysis pipeline for the behavioural and fMRI analyses. S.M.L and V.B.M collected single-cell data. R.L.B, G.d.H. and C.C.R. wrote and revised the manuscript, with input from S.M.L, V.B.M, M.G. and R.P.

## REFERENCES

1 Chib, V. S., Rangel, A., Shimojo, S. & O’Doherty, J. P. Evidence for a common representation of decision values for dissimilar goods in human ventromedial prefrontal cortex. Journal of Neuroscience 29, 12315–12320 (2009).

2 FitzGerald, T. H., Seymour, B. & Dolan, R. J. The role of human orbitofrontal cortex in value comparison for incommensurable objects. Journal of Neuroscience 29, 8388–8395 (2009).

3 Kable, J. W. & Glimcher, P. W. The neurobiology of decision: consensus and controversy. Neuron 63, 733–745 (2009).

4 Lebreton, M., Jorge, S., Michel, V., Thirion, B. & Pessiglione, M. An automatic valuation system in the human brain: evidence from functional neuroimaging. Neuron 64, 431–439 (2009).

5 Rangel, A., Camerer, C. & Montague, P. R. A framework for studying the neurobiology of value-based decision making. Nature reviews neuroscience 9, 545–556 (2008).

6 Rangel, A. & Hare, T. Neural computations associated with goal-directed choice. Curr Opin Neurobiol 20, 262–270 (2010).

7 Bartra, O., McGuire, J. T. & Kable, J. W. The valuation system: a coordinate-based meta-analysis of BOLD fMRI experiments examining neural correlates of subjective value. Neuroimage 76, 412–427 (2013).

8 Clithero, J. A. & Rangel, A. Informatic parcellation of the network involved in the computation of subjective value. Soc Cogn Affect Neurosci 9, 1289–1302 (2014).

9 Suzuki, S., Cross, L. & O’Doherty, J. P. Elucidating the underlying components of food valuation in the human orbitofrontal cortex. Nature neuroscience 20, 1780–1786 (2017).

10 O’Doherty, J. P., Rutishauser, U. & Iigaya, K. The hierarchical construction of value. Current opinion in behavioral sciences 41, 71–77 (2021).

11 Pessiglione, M. & Daunizeau, J. Bridging across functional models: The OFC as a value-making neural network. Behavioral Neuroscience 135, 277 (2021).

12 Iigaya, K. et al. Neural mechanisms underlying the hierarchical construction of perceived aesthetic value. Nature Communications 14, 127 (2023).

13 Polania, R., Woodford, M. & Ruff, C. C. Efficient coding of subjective value. Nature neuroscience 22, 134–142 (2019).

14 Rigoli, F. et al. A Bayesian model of context-sensitive value attribution. ELife 5, e16127 (2016).

15 Rigoli, F., Mathys, C., Friston, K. J. & Dolan, R. J. A unifying Bayesian account of contextual effects in value-based choice. PLoS computational biology 13, e1005769 (2017).

16 Ting, C.-C., Yu, C.-C., Maloney, L. T. & Wu, S.-W. Neural mechanisms for integrating prior knowledge and likelihood in value-based probabilistic inference. Journal of Neuroscience 35, 1792–1805 (2015).

17 Heng, J. A., Woodford, M. & Polania, R. Efficient sampling and noisy decisions. Elife 9, e54962 (2020).

18 Bouret, S. & Richmond, B. J. Ventromedial and orbital prefrontal neurons differentially encode internally and externally driven motivational values in monkeys. Journal of Neuroscience 30, 8591–8601 (2010).

19 Kennerley, S. W. & Wallis, J. D. Evaluating choices by single neurons in the frontal lobe: outcome value encoded across multiple decision variables. Eur J Neurosci 29, 2061–2073 (2009).

20 Padoa-Schioppa, C. & Assad, J. A. Neurons in the orbitofrontal cortex encode economic value. Nature 441, 223–226 (2006).

21 Roesch, M. R. & Olson, C. R. Neuronal activity related to reward value and motivation in primate frontal cortex. Science 304, 307–310 (2004).

22 Schoenbaum, G., Chiba, A. A. & Gallagher, M. Orbitofrontal cortex and basolateral amygdala encode expected outcomes during learning. Nature neuroscience 1, 155–159 (1998).

23 Tremblay, L. & Schultz, W. Reward-related neuronal activity during go-nogo task performance in primate orbitofrontal cortex. Journal of neurophysiology 83, 1864–1876 (2000).

24 Lopez-Persem, A. et al. Four core properties of the human brain valuation system demonstrated in intracranial signals. Nature Neuroscience 23, 664–675 (2020).

25 Rich, E. L. & Wallis, J. D. Decoding subjective decisions from orbitofrontal cortex. Nature neuroscience 19, 973–980 (2016).

26 Papageorgiou, G. K. et al. Inverted activity patterns in ventromedial prefrontal cortex during value-guided decision-making in a less-is-more task. Nature communications 8, 1886 (2017).

27 Kennerley, S. W. & Wallis, J. D. Encoding of reward and space during a working memory task in the orbitofrontal cortex and anterior cingulate sulcus. Journal of neurophysiology 102, 3352–3364 (2009).

28 Morrison, S. E. & Salzman, C. D. The convergence of information about rewarding and aversive stimuli in single neurons. Journal of Neuroscience 29, 11471–11483 (2009).

29 Padoa-Schioppa, C. Neuronal origins of choice variability in economic decisions. Neuron 80, 1322–1336 (2013).

30 Jazayeri, M. & Movshon, J. A. Optimal representation of sensory information by neural populations. Nature neuroscience 9, 690–696 (2006).

31 Ma, W. J., Beck, J. M., Latham, P. E. & Pouget, A. Bayesian inference with probabilistic population codes. Nature neuroscience 9, 1432–1438 (2006).

32 Pouget, A., Beck, J. M., Ma, W. J. & Latham, P. E. Probabilistic brains: knowns and unknowns. Nature neuroscience 16, 1170–1178 (2013).

33 Vilares, I. & Kording, K. Bayesian models: the structure of the world, uncertainty, behavior, and the brain. Annals of the New York Academy of Sciences 1224, 22–39 (2011).

34 Barretto-García, M. et al. Individual risk attitudes arise from noise in neurocognitive magnitude representations. Nature Human Behaviour 7, 1551–1567 (2023).

35 Chetverikov, A. & Jehee, J. F. M. Motion direction is represented as a bimodal probability distribution in the human visual cortex. Nature Communications 14, 7634 (2023).

36 Li, H.-H., Sprague, T. C., Yoo, A. H., Ma, W. J. & Curtis, C. E. Joint representation of working memory and uncertainty in human cortex. Neuron 109, 3699–3712. e3696 (2021).

37 Van Bergen, R. S. & Jehee, J. F. Probabilistic representation in human visual cortex reflects uncertainty in serial decisions. Journal of Neuroscience 39, 8164–8176 (2019).

38 Van Bergen, R. S., Ji Ma, W., Pratte, M. S. & Jehee, J. F. Sensory uncertainty decoded from visual cortex predicts behavior. Nature neuroscience 18, 1728–1730 (2015).

39 Walker, E. Y., Cotton, R. J., Ma, W. J. & Tolias, A. S. A neural basis of probabilistic computation in visual cortex. Nature Neuroscience 23, 122–129 (2020).

40 Zemel, R. S., Dayan, P. & Pouget, A. Probabilistic interpretation of population codes. Neural computation 10, 403–430 (1998).

41 Lange, R. D., Shivkumar, S., Chattoraj, A. & Haefner, R. M. Bayesian encoding and decoding as distinct perspectives on neural coding. Nature Neuroscience 26, 2063–2072 (2023).

42 Hangya, B., Sanders, J. I. & Kepecs, A. A mathematical framework for statistical decision confidence. Neural Computation 28, 1840–1858 (2016).

43 Meyniel, F., Sigman, M. & Mainen, Z. F. Confidence as Bayesian probability: From neural origins to behavior. Neuron 88, 78–92 (2015).

44 Pouget, A., Drugowitsch, J. & Kepecs, A. Confidence and certainty: distinct probabilistic quantities for different goals. Nature neuroscience 19, 366–374 (2016).

45 Geurts, L. S., Cooke, J. R., van Bergen, R. S. & Jehee, J. F. Subjective confidence reflects representation of Bayesian probability in cortex. Nature Human Behaviour 6, 294–305 (2022).

46 De Hollander, G., Grueschow, M., Hennel, F. & Ruff, C. C. Rapid Changes in Risk Attitudes Originate from Bayesian Inference on Parietal Magnitude Representations. BioRxiv, 2024.2008. 2023.609296 (2024).

47 Douglas, R. J. & Martin, K. A. Neuronal circuits of the neocortex. Annu. Rev. Neurosci. 27, 419–451 (2004).

48 Lebreton, M., Abitbol, R., Daunizeau, J. & Pessiglione, M. Automatic integration of confidence in the brain valuation signal. Nature neuroscience 18, 1159–1167 (2015).

49 Bedi, S., de Hollander, G. & Ruff, C. C. Probability weighting arises from boundary repulsions of cognitive noise. bioRxiv, 2025.2009. 2011.675565 (2025).

50 Grueschow, M., Polania, R., Hare, T. A. & Ruff, C. C. Automatic versus choice-dependent value representations in the human brain. Neuron 85, 874–885 (2015).

51 Anderson, B. A., Laurent, P. A. & Yantis, S. Value-driven attentional capture. Proceedings of the National Academy of Sciences 108, 10367–10371 (2011).

52 Pearson, D., Watson, P., Albertella, L. & Le Pelley, M. E. Attentional economics links value-modulated attentional capture and decision-making. Nature Reviews Psychology 1, 320–333 (2022).

53 Polania, R., Burdakov, D. & Hare, T. A. Rationality, preferences, and emotions with biological constraints: it all starts from our senses. Trends in Cognitive Sciences 28, 264–277 (2024).

54 Schaffner, J., Bao, S. D., Tobler, P. N., Hare, T. A. & Polania, R. Sensory perception relies on fitness-maximizing codes. Nature Human Behaviour 7, 1135–1151 (2023).

55 Deichmann, R., Gottfried, J. A., Hutton, C. & Turner, R. Optimized EPI for fMRI studies of the orbitofrontal cortex. Neuroimage 19, 430–441 (2003).

56 McGinty, V. B. & Lupkin, S. M. Behavioral read-out from population value signals in primate orbitofrontal cortex. Nat Neurosci 26, 2203–2212 (2023).

57 Kass, R. E. & Raftery, A. E. Bayes factors. Journal of the american statistical association 90, 773–795 (1995).

58 Bhatia, S. & Loomes, G. Noisy preferences in risky choice: A cautionary note. Psychological review 124, 678 (2017).

59 Abitbol, R. et al. Neural mechanisms underlying contextual dependency of subjective values: converging evidence from monkeys and humans. Journal of Neuroscience 35, 2308–2320 (2015).

60 Kurtz-David, V., Persitz, D., Webb, R. & Levy, D. J. The neural computation of inconsistent choice behavior. Nature communications 10, 1583 (2019).

61 Lee, J. K., Rouault, M. & Wyart, V. Adaptive tuning of human learning and choice variability to unexpected uncertainty. Science Advances 9, eadd0501 (2023).

62 Dabney, W. et al. A distributional code for value in dopamine-based reinforcement learning. Nature 577, 671–675 (2020).

63 Lowet, A. S., Zheng, Q., Matias, S., Drugowitsch, J. & Uchida, N. Distributional reinforcement learning in the brain. Trends in neurosciences 43, 980–997 (2020).

64 Gluth, S., Sommer, T., Rieskamp, J. & Büchel, C. Effective connectivity between hippocampus and ventromedial prefrontal cortex controls preferential choices from memory. Neuron 86, 1078–1090 (2015).

65 Renart, A. & Machens, C. K. Variability in neural activity and behavior. Current opinion in neurobiology 25, 211–220 (2014).

66 Gluth, S., Kern, N., Kortmann, M. & Vitali, C. L. Value-based attention but not divisive normalization influences decisions with multiple alternatives. Nature Human Behaviour 4, 634–645 (2020).

67 Krajbich, I. Accounting for attention in sequential sampling models of decision making. Current opinion in psychology 29, 6–11 (2019).

68 Rustichini, A., Domenech, P., Civai, C. & DeYoung, C. G. Working memory and attention in choice. Plos one 18, e0284127 (2023).

69 Olschewski, S., Rieskamp, J. & Scheibehenne, B. Taxing cognitive capacities reduces choice consistency rather than preference: A model-based test. Journal of Experimental Psychology: General 147, 462 (2018).

70 Lee, D. & Coricelli, G. An empirical test of the role of value certainty in decision making. Frontiers in psychology 11, 574473 (2020).

71 Cohen, J. D., McClure, S. M. & Yu, A. J. Should I stay or should I go? How the human brain manages the trade-off between exploitation and exploration. Philosophical Transactions of the Royal Society B: Biological Sciences 362, 933–942 (2007).

72 Daw, N. D., O’doherty, J. P., Dayan, P., Seymour, B. & Dolan, R. J. Cortical substrates for exploratory decisions in humans. Nature 441, 876–879 (2006).

73 Bertana, A., Chetverikov, A., van Bergen, R. S., Ling, S. & Jehee, J. F. Dual strategies in human confidence judgments. Journal of vision 21, 21–21 (2021).

74 Kepecs, A. & Mainen, Z. F. A computational framework for the study of confidence in humans and animals. Philosophical Transactions of the Royal Society B: Biological Sciences 367, 1322–1337 (2012).

75 De Martino, B., Fleming, S. M., Garrett, N. & Dolan, R. J. Confidence in value-based choice. Nature neuroscience 16, 105–110 (2013).

76 Lim, S.-L., O’Doherty, J. P. & Rangel, A. The decision value computations in the vmPFC and striatum use a relative value code that is guided by visual attention. Journal of Neuroscience 31, 13214–13223 (2011).

77 Boorman, E. D., Rushworth, M. F. & Behrens, T. E. Ventromedial prefrontal and anterior cingulate cortex adopt choice and default reference frames during sequential multi-alternative choice. Journal of neuroscience 33, 2242–2253 (2013).

78 Hunt, L. T. et al. Mechanisms underlying cortical activity during value-guided choice. Nature neuroscience 15, 470–476 (2012).

79 Albrecht, D. G., Geisler, W. S., Frazor, R. A. & Crane, A. M. Visual cortex neurons of monkeys and cats: temporal dynamics of the contrast response function. Journal of neurophysiology 88, 888–913 (2002).

80 Kayaert, G., Biederman, I., Op de Beeck, H. P. & Vogels, R. Tuning for shape dimensions in macaque inferior temporal cortex. Eur J Neurosci 22, 212–224 (2005).

81 Pruett Jr, J., Sinclair, R. & Burton, H. Response patterns in second somatosensory cortex (SII) of awake monkeys to passively applied tactile gratings. Journal of neurophysiology 84, 780–797 (2000).

82 Romo, R., Brody, C. D., Hernández, A. & Lemus, L. Neuronal correlates of parametric working memory in the prefrontal cortex. Nature 399, 470–473 (1999).

83 Zipser, D. & Andersen, R. A. A back-propagation programmed network that simulates response properties of a subset of posterior parietal neurons. Nature 331, 679–684 (1988).

84 Pouget, A. & Sejnowski, T. J. Spatial transformations in the parietal cortex using basis functions. Journal of cognitive neuroscience 9, 222–237 (1997).

85 Maunsell, J. H. & Van Essen, D. C. Functional properties of neurons in middle temporal visual area of the macaque monkey. I. Selectivity for stimulus direction, speed, and orientation. Journal of neurophysiology 49, 1127–1147 (1983).

86 Funahashi, S., Bruce, C. J. & Goldman-Rakic, P. S. Mnemonic coding of visual space in the monkey’s dorsolateral prefrontal cortex. Journal of neurophysiology 61, 331–349 (1989).

87 Logothetis, N. K., Pauls, J. & Poggio, T. Shape representation in the inferior temporal cortex of monkeys. Current biology 5, 552–563 (1995).

88 Georgopoulos, A. P., Kalaska, J. F., Caminiti, R. & Massey, J. T. On the relations between the direction of two-dimensional arm movements and cell discharge in primate motor cortex. Journal of Neuroscience 2, 1527–1537 (1982).

89 O’Keefe, J. & Dostrovsky, J. The hippocampus as a spatial map: preliminary evidence from unit activity in the freely-moving rat. Brain research (1971).

90 Enel, P., Wallis, J. D. & Rich, E. L. Stable and dynamic representations of value in the prefrontal cortex. Elife 9, e54313 (2020).

91 Hinkle, D. A. & Connor, C. E. Quantitative characterization of disparity tuning in ventral pathway area V4. Journal of neurophysiology 94, 2726–2737 (2005).

92 Zhang, T., Heuer, H. W. & Britten, K. H. Parietal area VIP neuronal responses to heading stimuli are encoded in head-centered coordinates. Neuron 42, 993–1001 (2004).

93 Peng, X. & Van Essen, D. C. Peaked encoding of relative luminance in macaque areas V1 and V2. Journal of neurophysiology 93, 1620–1632 (2005).

94 Lenninger, M., Skoglund, M., Herman, P. A. & Kumar, A. Are single-peaked tuning curves tuned for speed rather than accuracy? Elife 12, e84531 (2023).

95 Ghahramani, Z., Wolpert, D. M. & Jordan, M. I. Generalization to local remappings of the visuomotor coordinate transformation. Journal of Neuroscience 16, 7085–7096 (1996).

96 Guigon, E. Computing with populations of monotonically tuned neurons. Neural computation 15, 2115–2127 (2003).

97 Salinas, E. How behavioral constraints may determine optimal sensory representations. PLoS biology 4, e387 (2006).

98 Ecker, A., Berens, P., Tolias, A. & Bethge, M. The effect of noise correlations in populations of diversely tuned neurons. Nature Precedings, 1–1 (2011).

99 Zylberberg, J. The role of untuned neurons in sensory information coding. BioRxiv, 134379 (2017).

100 Esteban, O. et al. fMRIPrep: a robust preprocessing pipeline for functional MRI. Nature methods 16, 111–116 (2019).

