## Supplementary Information for "A Neural Population Code for Value in Human Orbitofrontal Cortex"

### Anatomical data preprocessing

The T1-weighted (T1w) image was corrected for intensity non-uniformity (INU) with N4BiasFieldCorrection (Tustison et al. 2010), distributed with ANTs 2.2.0 (Avants et al. 2008, RRID:SCR_004757), and used as T1w-reference throughout the workflow. The T1w-reference was then skull-stripped with a Nipype implementation of the antsBrainExtraction.sh workflow (from ANTs), using OASIS30ANTs as target template. Brain tissue segmentation of cerebrospinal fluid (CSF), white-matter (WM) and gray-matter (GM) was performed on the brain-extracted T1w using fast (FSL 5.0.9, RRID:SCR_002823, Zhang, Brady, and Smith 2001). Brain surfaces were reconstructed using recon-all (FreeSurfer 6.0.1, RRID:SCR_001847, Dale, Fischl, and Sereno 1999), and the brain mask estimated previously was refined with a custom variation of the method to reconcile ANTs-derived and FreeSurfer-derived segmentations of the cortical gray-matter of Mindboggle (RRID:SCR_002438, Klein et al. 2017). Volume-based spatial normalization to one standard space (MNI152NLin2009cAsym) was performed through nonlinear registration with antsRegistration (ANTs 2.2.0), using brain-extracted versions of both T1w reference and the T1w template. The following template was selected for spatial normalization: ICBM 152 Nonlinear Asymmetrical template version 2009c [Fonov et al. (2009), RRID:SCR_008796; TemplateFlow ID: MNI152NLin2009cAsym].

### Functional data preprocessing

For each of the 8 BOLD runs found per subject (across all tasks and sessions), the following preprocessing was performed. First, a reference volume and its skull-stripped version were generated using a custom methodology of fMRIPrep. The BOLD reference was then co-registered to the T1w reference using bbregister (FreeSurfer) which implements boundary-based registration (Greve and Fischl 2009). Co-registration was configured with six degrees of freedom. Head-motion parameters with respect to the BOLD reference (transformation matrices, and six corresponding rotation and translation parameters) are estimated before any spatiotemporal filtering using mcflirt (FSL 5.0.9, Jenkinson et al. 2002). BOLD runs were slice-time corrected using 3dTshift from AFNI 20160207 (Cox and Hyde 1997, RRID:SCR_005927). The BOLD time-series, were resampled to surfaces on the following spaces: fsaverage5, fsaverage6. The BOLD time-series (including slice-timing correction when applied) were resampled onto their original, native space by applying a single, composite transform to correct for head-motion and susceptibility distortions. These resampled BOLD time-series will be referred to as preprocessed BOLD in original space, or just preprocessed BOLD. The BOLD time-series were resampled into standard space, generating a preprocessed BOLD run in [‘MNI152NLin2009cAsym’] space. First, a reference volume and its skull-stripped version were generated using a custom methodology of fMRIPrep. Several confounding time-series were calculated based on the preprocessed BOLD: framewise displacement (FD), DVARS and three region-wise global signals. FD and DVARS are calculated for each functional run, both using their implementations in Nipype (following the definitions by Power et al. 2014). The three global signals are extracted within the CSF, the WM, and the whole-brain masks. Additionally, a set of physiological regressors were extracted to allow for component-based noise correction (CompCor, Behzadi et al. 2007). Principal components are estimated after high-pass filtering the preprocessed BOLD time-series (using a discrete cosine filter with 128s cut-off) for the two CompCor variants: temporal (tCompCor) and anatomical (aCompCor). tCompCor components are then calculated from the top 5% variable voxels within a mask covering the subcortical regions. This subcortical mask is obtained by heavily eroding the brain mask, which ensures it does not include cortical GM regions. For aCompCor, components are calculated within the intersection of the aforementioned mask and the union of CSF and WM masks calculated in T1w space, after their projection to the native space of each functional run (using the inverse BOLD-to-T1w transformation). Components are also calculated separately within the WM and CSF masks. For each CompCor decomposition, the k components with the largest singular values are retained, such that the retained components’ time series are sufficient to explain 50 percent of variance across the nuisance mask (CSF, WM, combined, or temporal). The remaining components are dropped from consideration. The head-motion estimates calculated in the correction step were also placed within the corresponding confounds file. The confound time series derived from head motion estimates and global signals were expanded with the inclusion of temporal derivatives and quadratic terms for each (Satterthwaite et al. 2013). Frames that exceeded a threshold of 0.5 mm FD or 1.5 standardised DVARS were annotated as motion outliers. All resamplings can be performed with a single interpolation step by composing all the pertinent transformations (i.e. head-motion transform matrices, susceptibility distortion correction when available, and co-registrations to anatomical and output spaces). Gridded (volumetric) resamplings were performed using antsApplyTransforms (ANTs), configured with Lanczos interpolation to minimize the smoothing effects of other kernels (Lanczos 1964). Non-gridded (surface) resamplings were performed using mri_vol2surf (FreeSurfer).

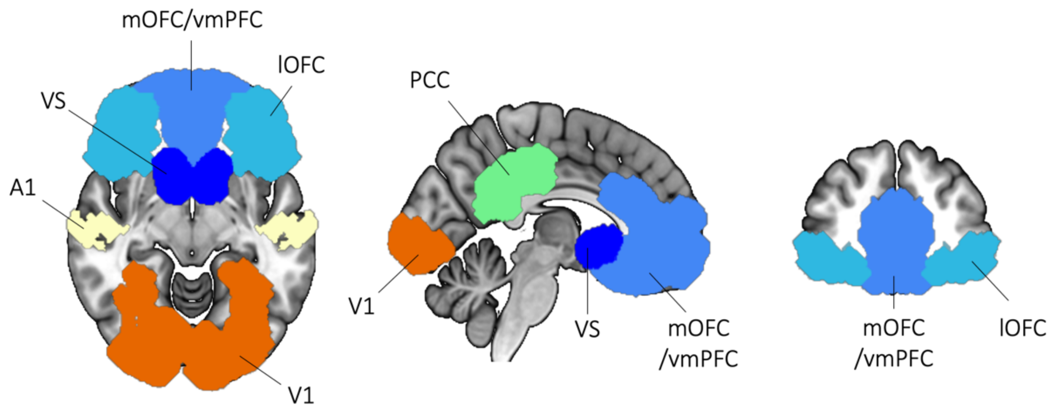


**Supplementary Fig. 1. Brain regions of interest (ROIs).** ROIs were anatomically defined using the MarsAtlas-Colin27-MNI cortical parcellation atlas (cortical structures) and the Harvard-Oxford atlas (subcortical structures; see Methods). mOFC: medial orbitofrontal cortex; vmPFC: ventromedial prefrontal cortex; lOFC: lateral orbitofrontal cortex; VS: ventral striatum; A1: primary auditory cortex; V1: primary visual cortex; PCC: posterior cingulate cortex.

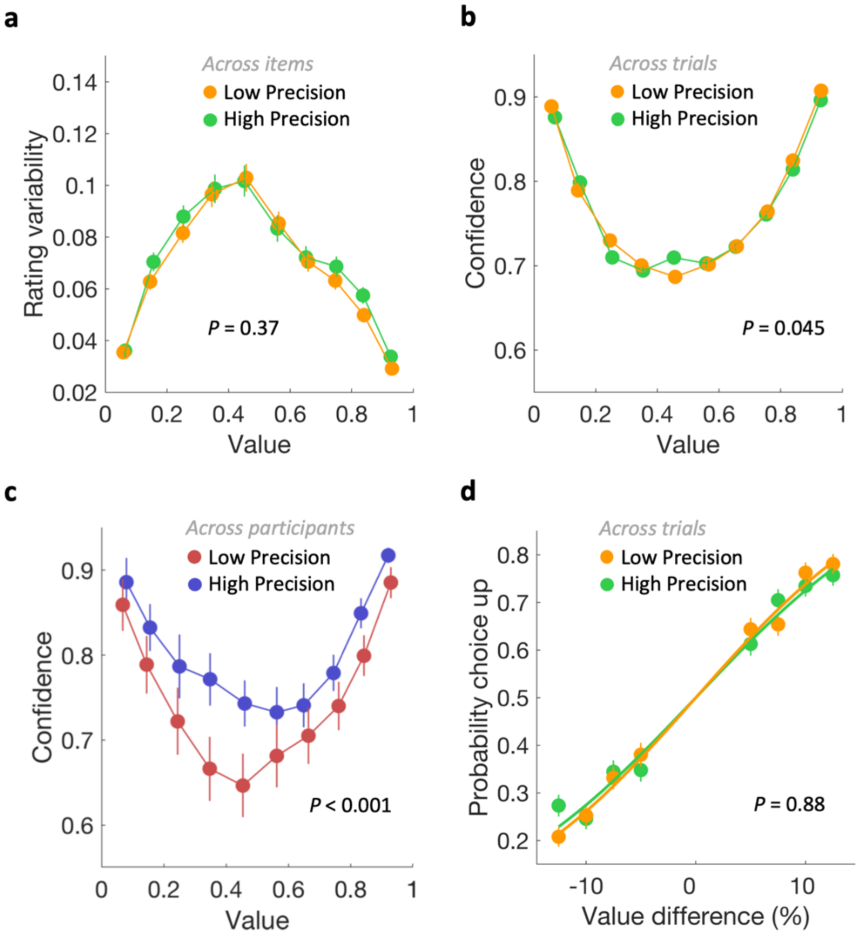


**Supplementary Fig. 2. Relationships between precision of neural value representation in the lateral OFC and behavioural variability. a,** value rating variability plotted as a function of each item's mean rating (in 10 value bins) across both rounds for experiment 1. Dots and error bars represent the mean and s.e.m. across items, with low (orange) or high (green) decoded neural precision. **b,c,** confidence ratings plotted as a function of each item's mean rating (in 10 value bins) averaged across both rounds for experiment 2. Dots and error bars represent the mean and s.e.m. across trials (**b**) with low (orange) or high (green) decoded neural precision; and across participants (**c**) with low (red) or high (blue) averaged decoded neural precision. **d,** Observed choice probability plotted as a function of the two item's value difference ($V_1 - V_2$, in 8 bins) for experiment 1. Dots and error bars represent the mean and s.e.m. across trials (**d**) with low (orange) or high (green) decoded neural precision.

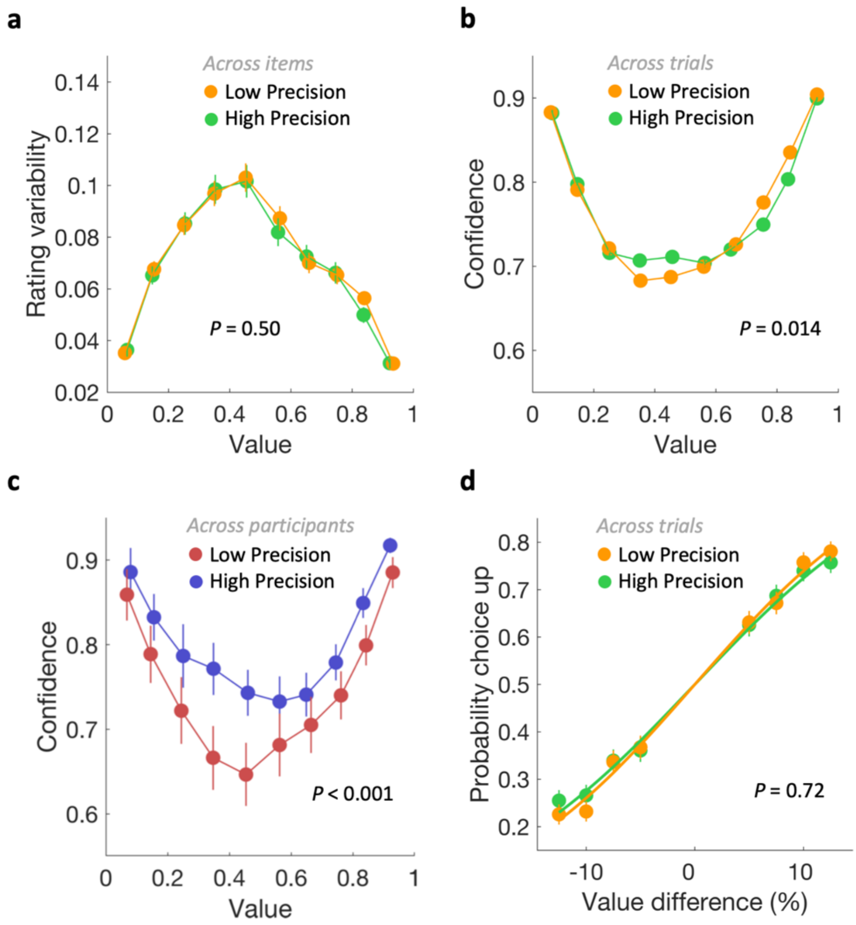


**Supplementary Fig. 3. Relationships between precision of neural value representation in the PCC and behavioural variability. a,** value rating variability plotted as a function of each item's mean rating (in 10 value bins) across both rounds for experiment 1. Dots and error bars represent the mean and s.e.m. across items, with low (orange) or high (green) decoded neural precision. **b,c,** confidence ratings plotted as a function of each item's mean rating (in 10 value bins) averaged across both rounds for experiment 2. Dots and error bars represent the mean and s.e.m. across trials (**b**) with low (orange) or high (green) decoded neural precision; and across participants (**c**) with low (red) or high (blue) averaged decoded neural precision. **d,** Observed choice probability plotted as a function of the two item's value difference ($V_1 - V_2$, in 8 bins) for experiment 1. Dots and error bars represent the mean and s.e.m. across trials (**d**) with low (orange) or high (green) decoded neural precision.

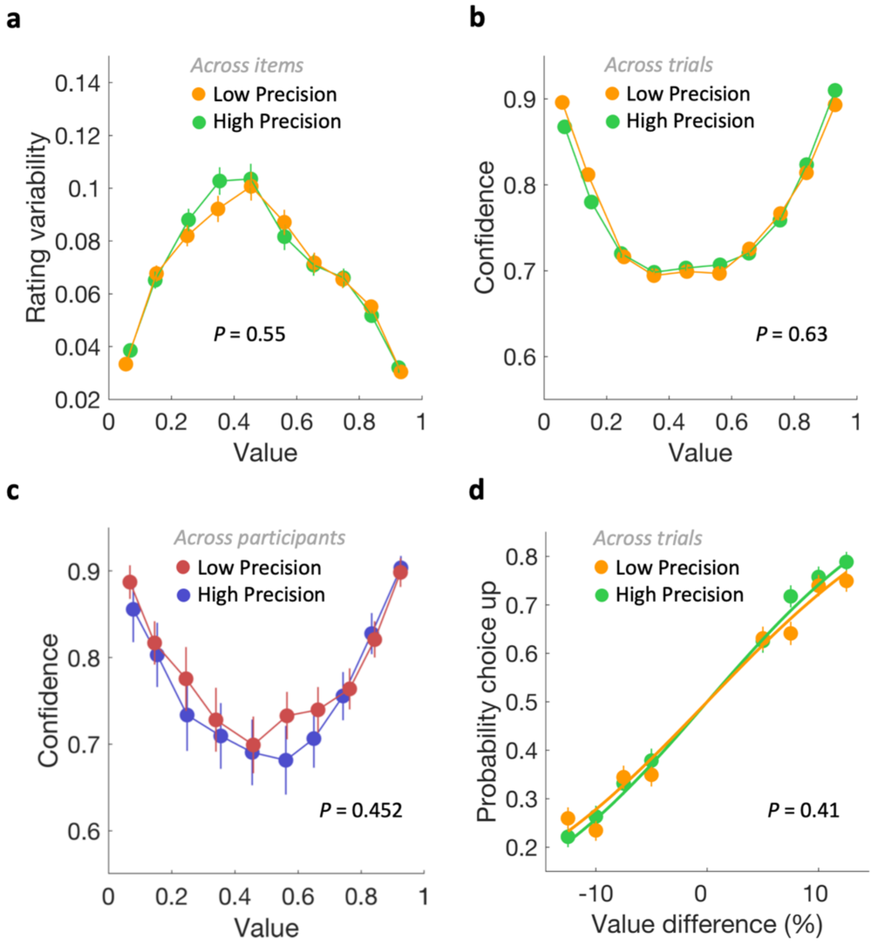


**Supplementary Fig. 4. Relationships between precision of neural value representation in the ventral striatum and behavioural variability. a,** value rating variability plotted as a function of each item's mean rating (in 10 value bins) across both rounds for experiment 1. Dots and error bars represent the mean and s.e.m. across items, with low (orange) or high (green) decoded neural precision. **b,c,** confidence ratings plotted as a function of each item's mean rating (in 10 value bins) averaged across both rounds for experiment 2. Dots and error bars represent the mean and s.e.m. across trials (**b**) with low (orange) or high (green) decoded neural precision; and across participants (**c**) with low (red) or high (blue) averaged decoded neural precision. **d,** Observed choice probability plotted as a function of the two item's value difference ($V_1 - V_2$, in 8 bins) for experiment 1. Dots and error bars represent the mean and s.e.m. across trials (**d**) with low (orange) or high (green) decoded neural precision.

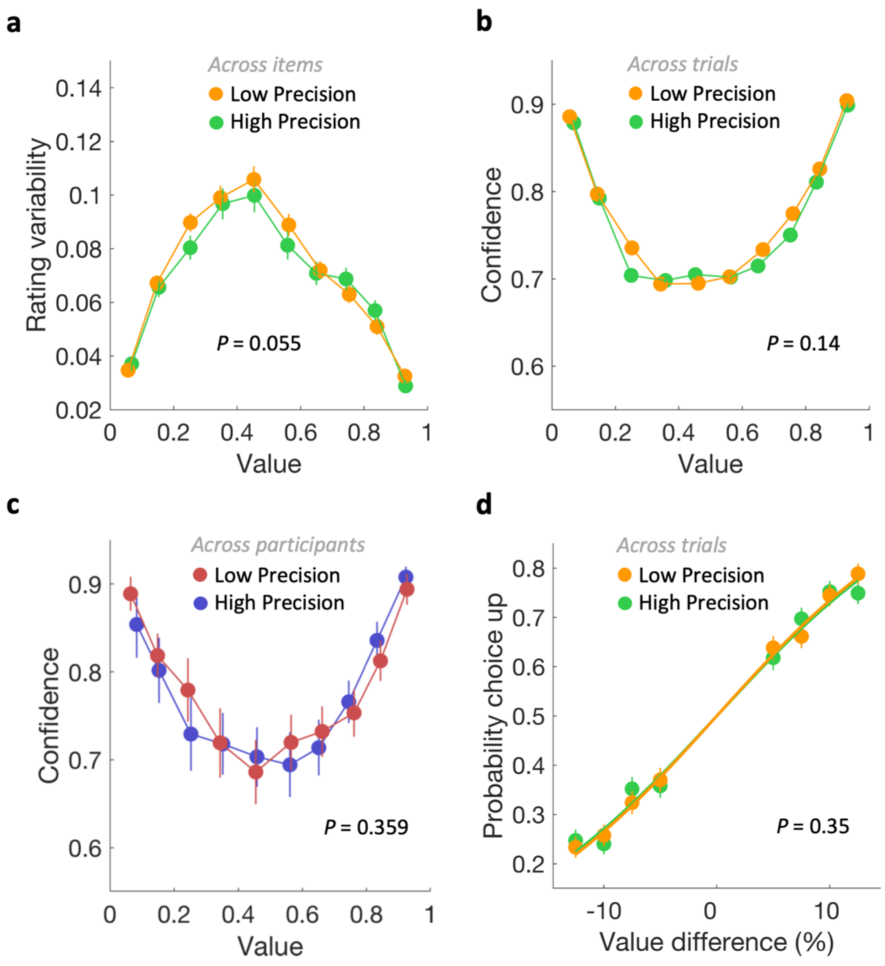


**Supplementary Fig. 5. Relationships between precision of neural value representation in the primary visual cortex and behavioural variability. a,** value rating variability plotted as a function of each item's mean rating (in 10 value bins) across both rounds for experiment 1. Dots and error bars represent the mean and s.e.m. across items, with low (orange) or high (green) decoded neural precision. **b,c,** confidence ratings plotted as a function of each item's mean rating (in 10 value bins) averaged across both rounds for experiment 2. Dots and error bars represent the mean and s.e.m. across trials (**b**) with low (orange) or high (green) decoded neural precision; and across participants (**c**) with low (red) or high (blue) averaged decoded neural precision. **d,** Observed choice probability plotted as a function of the two item's value difference ($V_1 - V_2$, in 8 bins) for experiment 1. Dots and error bars represent the mean and s.e.m. across trials (**d**) with low (orange) or high (green) decoded neural precision.

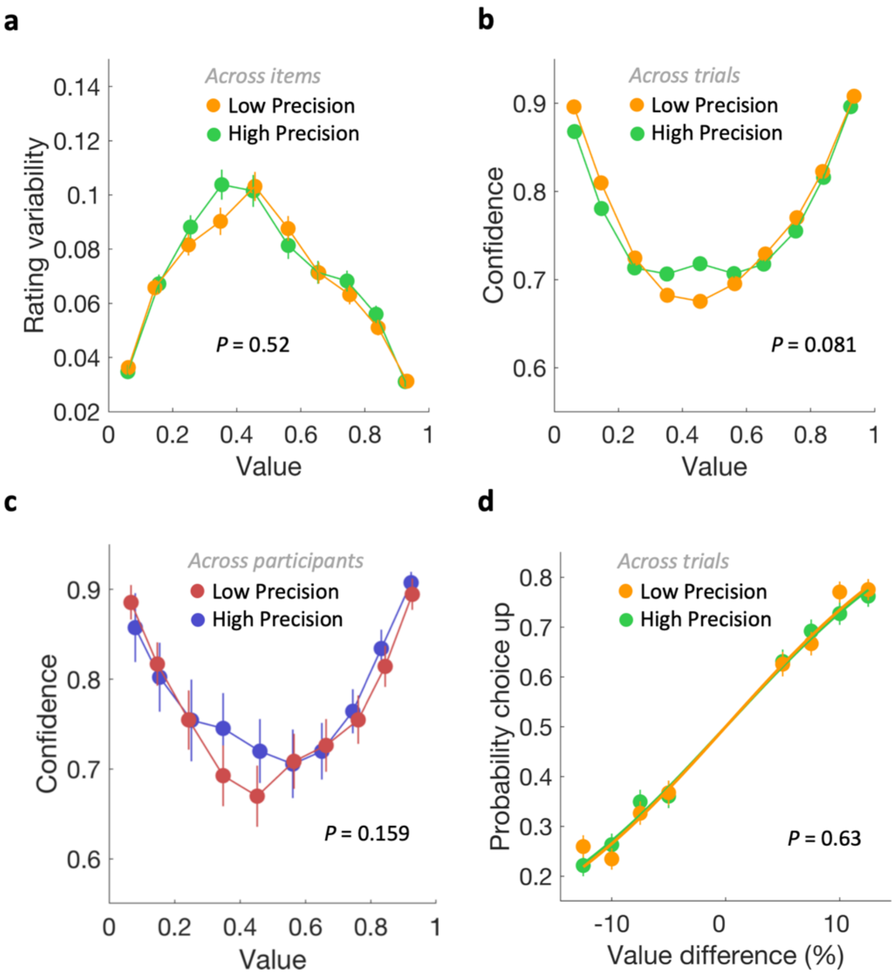


**Supplementary Fig. 6. Relationships between precision of neural value representation in the auditory cortex and behavioural variability. a,** value rating variability plotted as a function of each item's mean rating (in 10 value bins) across both rounds for experiment 1. Dots and error bars represent the mean and s.e.m. across items, with low (orange) or high (green) decoded neural precision. **b,c,** confidence ratings plotted as a function of each item's mean rating (in 10 value bins) averaged across both rounds for experiment 2. Dots and error bars represent the mean and s.e.m. across trials (**b**) with low (orange) or high (green) decoded neural precision; and across participants (**c**) with low (red) or high (blue) averaged decoded neural precision. **d,** Observed choice probability plotted as a function of the two item's value difference ($V_1 - V_2$, in 8 bins) for experiment 1. Dots and error bars represent the mean and s.e.m. across trials (**d**) with low (orange) or high (green) decoded neural precision.

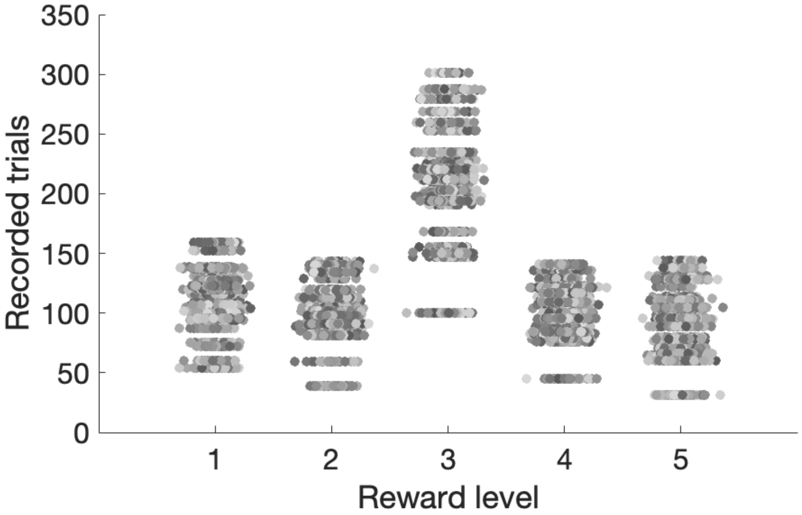


**Supplementary Fig. 7. Number of recorded trials for OFC cells.** Plots show the number of trials recorded for each of the 1,450 OFC neurons at each of the five value levels. Each dot represents a neuron. Neurons recorded using the same multi-channel linear recording arrays share identical trial counts. For each neuron, activity was recorded over 50–300 repetitions per value level, ensuring robust estimation of responses across value levels.
